# Sphingosine kinase 2 is essential for remyelination following cuprizone intoxication

**DOI:** 10.1101/2021.02.14.431183

**Authors:** Huitong Song, Holly P. McEwen, Thomas Duncan, Jun Yup Lee, Jonathan D. Teo, Anthony S. Don

## Abstract

Therapeutics that promote oligodendrocyte survival and remyelination are needed to restore neurological function in demyelinating diseases. Sphingosine 1-phosphate (S1P) is an essential lipid metabolite that signals through five G-protein coupled receptors. S1P receptor agonists such as Fingolimod are valuable immunosuppressants used to treat multiple sclerosis, and promote oligodendrocyte survival. However, the role for endogenous S1P, synthesized by the enzyme sphingosine kinase 2 (SphK2), in oligodendrocyte survival and myelination has not been established. This study investigated the requirement for SphK2 in oligodendrocyte survival and remyelination using the cuprizone mouse model of acute demyelination, followed by spontaneous remyelination. Oligodendrocyte density did not differ between untreated wild-type (WT) and SphK2 knockout (SphK2^-/-^) mice. However, cuprizone treatment caused significantly greater loss of mature oligodendrocytes in SphK2^-/-^ compared to WT mice. Following cuprizone withdrawal, spontaneous remyelination occurred in WT but not SphK2^-/-^ mice, even though progenitor and mature oligodendrocyte density increased in both genotypes. Levels of cytotoxic sphingosine and ceramide were higher in the corpus callosum of SphK2^-/-^ mice, and in contrast to WT mice, did not decline following cuprizone withdrawal in SphK2^-/-^ mice. We also observed a significant reduction in myelin thickness with ageing in SphK2^-/-^ compared to WT mice. These results provide the first evidence that SphK2, the dominant enzyme catalysing S1P synthesis in the adult brain, is essential for remyelination following a demyelinating insult and myelin maintenance with ageing. We propose that persistently high levels of sphingosine and ceramide, a direct consequence of SphK2 deficiency, may block remyelination.

**Table of Contents Image:** 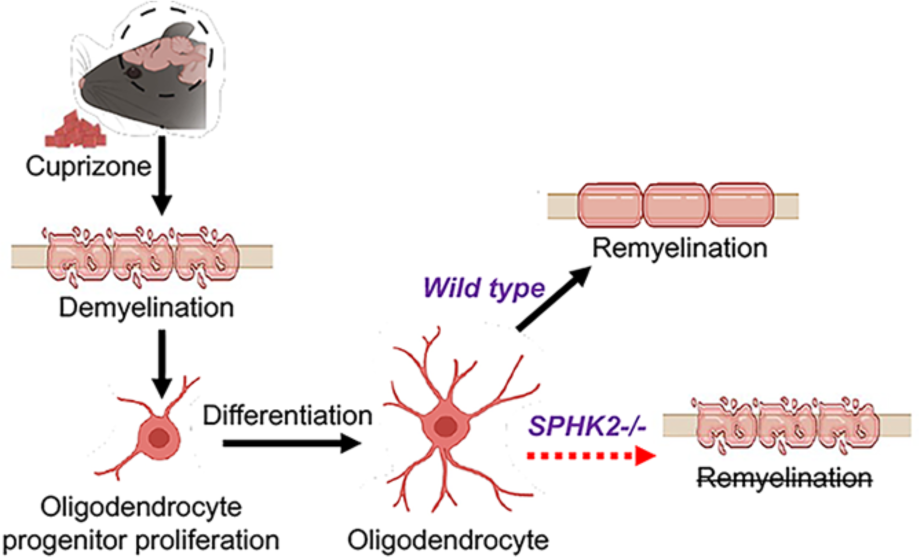

**Main Points:**

- The lipid kinase sphingosine kinase 2 (SphK2) is essential for remyelination following a demyelinating insult (cuprizone).
- SphK2 protects against cuprizone-mediated oligodendrocyte loss.
- SphK2 deficiency leads to thinner myelin with ageing.

## Introduction

Myelin is a multi-layered membrane that surrounds neuronal axons and is necessary for the rapid and energy-efficient propagation of action potentials (Roth & Nunez, 2016; Simons & Nave, 2016). Myelin is synthesized by oligodendrocytes in the central nervous system (CNS) and Schwann cells in the peripheral nervous system. Oligodendrocytes and myelin are essential not only for rapid impulse propagation, but also for neuronal bioenergetics and survival (Simons & Nave, 2016).

Immune-mediated degradation of myelin is a pathological hallmark of multiple sclerosis (MS) (Reich et al., 2018). This inflammatory demyelination is believed to cause loss of neurological function through reduced efficiency of axonal conduction, loss of oligodendrocyte support for axons, and direct effects of local inflammation on axons (Plemel et al., 2017; Reich et al., 2018). Immunosuppressive therapies are generally effective at blunting neuroinflammation and demyelination in relapsing-remitting MS, which is the most common form of the disease. This slows disease progression, however endogenous myelin repair is limited. Relapsing-remitting MS eventually progresses into secondary progressive MS, characterised by irreversible decline. Therapeutics that promote oligodendrocyte survival and remyelination are needed to restore neurological function in MS and other demyelinating diseases (Plemel et al., 2017; Stangel et al., 2017).

Myelin is comprised 70-80% of lipids, and is enriched in the sphingolipids galactosylceramide (a.k.a. cerebroside), sulfatide, and sphingomyelin, which are all synthesized from ceramide (Schmitt et al., 2015). Alternatively, ceramide can be de-acylated by ceramidases to produce sphingosine, which is phosphorylated by sphingosine kinase 1 (SphK1) or 2 (SphK2) to produce the signalling lipid S1P (Grassi et al., 2019). SphK2 is the dominant isoform in the brain, as S1P levels are ∼85% lower in grey matter of SphK2 knockout (SphK2^-/-^) mice (Lei et al., 2017). In accordance with the high levels of sphingolipids in myelin, S1P levels are 3- to 10-fold higher in white matter compared to grey matter (Couttas et al., 2014; Wang et al., 2019).

After its synthesis, S1P is secreted and signals through a family of five G-protein coupled receptors, S1PR1-S1PR5 (Choi & Chun, 2013; Grassi et al., 2019). S1P receptors 1-3 (S1PR1- 3) are expressed by most cells and tissues, although S1PR1 expression is particularly high in astrocytes and endothelial cells. S1PR4 expression is restricted to immune cells, whilst S1PR5 is expressed almost exclusively by mature oligodendrocytes in the CNS (Jaillard et al., 2005; Y. Zhang et al., 2014). *In vitro* cell culture studies have shown that S1P, signalling through S1PR1, stimulates oligodendrocyte progenitor cell (OPC) proliferation (Jung et al., 2007), whereas S1PR5 promotes oligodendrocyte survival (Jaillard et al., 2005).

The MS drug Fingolimod is a sphingosine analogue that, when phosphorylated by SphK2 *in vivo*, becomes a potent agonist of S1P receptors 1, 3, 4, and 5 (Bigaud et al., 2014; Mandala et al., 2002). Its clinical success led to the recent approval of selective S1PR1/S1PR5 agonists Siponimod and Ozanimod for treatment of MS, with other S1PR1 agonists currently in clinical trials (Bigaud et al., 2014; Stepanovska & Huwiler, 2020). Suppression of autoimmunity with this class of drugs is attributed primarily to hyper-activation of S1PR1 on lymph-node resident lymphocytes. This impedes the S1PR1-dependent egress of lymphocytes from lymph nodes into the circulation (Mandala et al., 2002; Matloubian et al., 2004), and is therefore termed “functional antagonism” of S1PR1. In addition to its peripheral effects, fingolimod readily crosses the blood brain barrier, and the effectiveness of the drug in experimental autoimmune encephalitis (EAE), a rodent model of autoimmune demyelination, is at-least partly attributed to its action on astrocyte S1PR1 (Choi et al., 2011). Fingolimod also protects against demyelination induced with the oligodendrocyte toxin cuprizone (H. J. Kim et al., 2011; S. Kim et al., 2018), but does not enhance spontaneous remyelination after cuprizone withdrawal (Alme et al., 2015; Nystad et al., 2020). It remains to be established if endogenous S1P is an important mediator of oligodendrocyte survival and remyelination *in vivo*.

We recently demonstrated that SphK2 deficiency synergises with an amyloidogenic transgene to cause oligodendrocyte loss and demyelination in a mouse model of Alzheimer’s disease (Lei et al., 2019). Herein we employed the oligodendrocyte toxin cuprizone to determine if endogenous SphK2 protects against demyelination and/or is necessary for remyelination following a well-established demyelinating insult. Cuprizone administration in the diet is accompanied by acute demyelination attributed to oligodendrocyte death (Benardais et al., 2013; Zhan et al., 2020). Spontaneous remyelination occurs within 2-3 weeks of cuprizone withdrawal, making this an ideal model for investigating factors that regulate oligodendrocyte survival and myelin elaboration in adult mice. We demonstrate that endogenous SphK2 protects against cuprizone-mediated oligodendrocyte loss, and is essential for remyelination following cuprizone withdrawal. We also show that myelin sheaths are thinner in aged SphK2^- /-^ mice, suggesting that SphK2 is necessary for myelin turnover or biosynthesis in the adult mouse, but not for developmental myelination.

## Experimental Methods

### Transgenic mice and cuprizone administration

SphK2 knockout mice (SphK2^−/−^) (RRID: IMSR_JAX:019140) were obtained under MTA with Prof Richard Proia, NIDDK, Bethesda, USA and genotyped as described (Mizugishi et al., 2005). SphK2^−/−^ mice were crossed to C57BL/6J mice for more than 10 generations. SphK2^−/−^ and wild-type (WT) control (SphK2^+/+^) mice used in this project were littermates derived from SphK2^+/−^ breeding pairs. Male mice were used for all experiments. Mice were housed in independently-ventilated cages, with food and water provided *ad libitum* and a 12 h light/dark schedule, and weighed twice per week. Experiments were conducted in accordance with the Australian Code of Practice for the Care and Use of Animals for Scientific Purposes, and approved by the University of Sydney animal ethics committee (2017/1284).

Groups of 5-8 mice, at 12 weeks of age, were randomly allocated to treated (0.2% (w/w) cuprizone in normal chow) or untreated (normal chow) groups. Cuprizone (Sigma Aldrich, #C9012) was incorporated into chow pellets by Specialty Feeds, Australia. Mice were culled by isofluorane inhalation, then perfused transcardially with sterile saline. Brains were removed and halved sagittally. One hemisphere was post-fixed overnight with 4% paraformaldehyde (Sigma-Aldrich, #28908) in PBS, then transferred to 30% sucrose and frozen at −80 °C. Frozen tissue was sectioned at 30 μm using a ThermoFisher Scientific Cyrotome FSE Cryostat and stored at 30 °C in cryoprotectant (25% glycerol, 25% ethylene glycol in PBS). Corpus callosum (CC), cortex, and hippocampus were dissected from the other hemisphere and flash-frozen in liquid nitrogen for biochemical analyses.

### Luxol fast blue/cresyl violet staining

Sections were mounted on Superfrost Plus microscope slides (Thermofisher Scientific #MENSF41296SP) and dried at 37 °C for 3 h. Sections were defatted for 5 h in 1:1 alcohol/chloroform, soaked in 95% ethanol for 30 min, then incubated for 16 h at 60 °C in 1 mg/ml luxol fast blue (Sigma-Aldrich, #S3382) in 95% ethanol. Sections were rinsed with 80% ethanol followed by water, differentiated in 0.05% Li2CO3 (Sigma-Aldrich #255823) solution for 30 min, rinsed in 70% ethanol and water, then counterstained with cresyl violet (1 mg/mL cresyl violet and 0.05 mg/mL oxalic acid), and rinsed successively in water, 70% ethanol, 95% ethanol, 100% ethanol and xylene. DPX mountant (Sigma-Aldrich #06522) was used for mounting, and slides were dried overnight. Images were acquired using a Zeiss Axio Scan.Z1. A semiquantitative scoring system (0-3) (Doan et al., 2013; Gudi et al., 2009) was used to quantify myelin, with a score of 3 indicating intact myelin and 0 indicating complete demyelination. Myelin integrity in the splenium, body and genu of the CC was scored by three blinded investigators (independently). These scores were averaged to provide the myelination score for each mouse.

### Immunofluorescence staining

For ASPA, GFAP and Iba1 staining, free-floating sections were incubated for 10 min at 70 °C in 10 mM sodium citrate, pH 6.0, 0.01% Tween 20, followed by 3 washes with PBS containing 0.1% Tween 20 (PBST). Sections were then incubated in blocking solution (5% goat serum, 0.1% BSA, 0.1% Triton X-100 in PBS) for 2 h at RT, and overnight at 4 °C in primary antibodies (Table 1) diluted in blocking solution. One section was incubated in blocking buffer without primary antibody as a control for non-specific binding of secondary antibodies. Sections were then washed and incubated with AlexaFluor 488-, 546-, or 647-conjugated secondary antibodies (1:500 in blocking solution) for 2 h at RT in the dark. After 3 washes with PBST, sections were stained with 1 µg/mL diamidino-2-phenylindole dihydrochloride (DAPI) for 7 min, washed 2 times with PBST, and mounted on poly-L-Lysine coated slides (ProSciTec #G312P-W) using ProLong Glass anti-fade (Life Technologies #P36980). For NG2/Olig2 staining, permeabilization was performed using 0.3% Triton X-100 in PBS for 10 min, blocking solution was 5% BSA, 0.1% Triton X-100 in PBS, primary antibodies were diluted in PBS containing 2.5% BSA and 0.05% Triton X-100, and sections were incubated with donkey anti-goat Alexa-488 and goat anti-rabbit Alexa-546 antibodies sequentially for 2 h each time, with four washes in between.

**Table 1.** Primary antibodies used for immunohistochemistry (IHC) and western blotting (WB).

| Primary antibody | Supplier | Dilution | Cat. # | RRID |
| --- | --- | --- | --- | --- |
| Rabbit anti-ASPA | Abcam | 1:1000<br>(WB/IHC) | ab223269 | None found |
| Rabbit anti-CerS2 | Thermo Fisher | 1:1000 (WB) | PA5-55560 | AB_2643295 |
| Rabbit anti-MBP | Abcam | 1:1000<br>(WB/IHC) | ab40390 | AB_1141521 |
| Rabbit anti-MOG | Abcam | 1:1000 (WB) | ab32760 | AB_2145529 |
| Rabbit anti-PLP | Abcam | 1:1000 (WB) | ab28486 | AB_776593 |
| Mouse anti-CNP [11-5B] | Abcam | 1:1000 (WB) | ab6319 | AB_2082593 |
| Rabbit anti-β-actin | Abcam | 1:5000 (WB) | ab8227 | AB_2305186 |
| Rabbit anti-NG2 | Sigma Aldrich | 1:500 (IHC) | ZRB5320 | None found |
| Goat anti-Olig2 | R&D Systems | 1:500 (IHC) | AF2418 | AB_2157554 |
| Rabbit Anti-Iba1 | Abcam | 1:1000 (IHC) | ab178847 | AB_2832244 |
| Mouse anti-GFAP (GA5) | Cell Signalling Technology | 1:1000 (IHC) | 3670S | AB_561049 |
| Mouse anti-S1PR1 (A6) | Santa Cruz Biotechnology | 1:500 (WB) | sc-48356 | AB_2238920 |
| Rabbit anti-S1PR5 | Novus Biologicals | 1:500 (WB) | NBP2-24712SS | AB_2888942 |

ASPA-positive cells were imaged using an Olympus Virtual Slide microscope VS1200 and counted using the particle analysis function of ImageJ (v. 1.53b), based on size (7.50-100.00 μm^2^) and circularity (0.25-1.00). NG2/Olig2, GFAP and Iba1 staining were imaged with a Leica SP8 microscope. NG2/Olig2 double-positive cells were counted manually by a blinded observer. Olig2-positive cells were counted by ImageJ using the same method as ASPA-positive cells. Cell density was calculated as cell number divided by CC area. GFAP and Iba1 staining were quantified as the area of CC occupied by the fluorescent stain, using ImageJ. In all cases, the experimental operator and analyser remained blinded to the treatment groups until after quantification.

### MBP immunohistochemistry

Free-floating sections were permeabilized in PBS with 0.3% Triton X-100 for 10 min, then washed three times in PBST. Sections were then blocked for 2 h at RT in 5% skim milk, 2% goat serum, 0.1% BSA, 0.1% Triton X-100 in PBS, and incubated overnight at 4 °C with rabbit anti-MBP diluted in 5% goat serum, 0.1% BSA, 0.1% Triton X-100 in PBS. After 3 washes with PBST, endogenous peroxidase was blocked by incubating for 10 min at RT in 0.3% H2O2 (Sigma-Aldrich #216763). Sections were then incubated in biotinylated goat anti-rabbit antibody (Abcam #ab6720, RRID: AB_954902, 1:500) for 2 h, washed three times, and incubated with horseradish peroxidase (HRP)-conjugated Streptavidin (Vector Laboratories #SA-5004-1, 1:500 in PBS) for 2 h. Slides were washed three times, and signal was developed with the DAB Peroxidase Substrate Kit (Vector Laboratories #SK-4100). Sections were then rinsed in PBST, mounted and dried. Slides were dehydrated and embedded as described for luxol fast blue staining. Images were acquired using a Zeiss Axio Scan.Z1. MBP staining in the CC was quantified using the same semi-quantitative scoring system as used for luxol fast blue staining.

### Western blotting

Tissue samples (10-20 mg) were homogenized at 4 °C in 0.4 mL radio-immunoprecipitation assay buffer (10 mM pH 7.4 Tris, 100 mM NaCl, 100 mM sodium pyrophosphate, 0.5% sodium deoxycholate, 1% Triton-X, 10% glycerol, 1 mM EDTA, 1 mM NaF, 0.1% SDS) supplemented with Complete Mini, EDTA-free Protease Inhibitor Cocktail (Sigma #11836170001) and phosphatase inhibitors (5 mM β-glycerophosphate, 2 mM sodium orthovanadate, and 5 mM NaF), using a Biospec mini bead beater with acid-washed glass beads (425-600 μm). Extracts were cleared by centrifugation at 1000 g for 15 min, at 4 °C, and protein concentration was measured by bicinchoninic acid assay (ThermoFisher Scientific #23225). Protein lysates were resolved on Bolt™ 4-12% Bis-Tris Plus gels (ThermoFisher Scientific #NW04125BOX) and transferred to polyvinylidene fluoride membranes. Membranes were blocked for 1 h with 5% skim milk in Tris-buffered saline containing 0.1% Tween 20 (TBST), then washed three times with TBST and incubated overnight (4 °C) with primary antibody (Table 1) in TBST containing 3% BSA. Membranes were washed three times with TBST, incubated with HRP-conjugated secondary antibody (Cell Signalling #7074, RRID: AB_2099233, or #7076, RRID: AB_330924) in blocking buffer for 1 h, and washed again before imaging. Signal was developed with ECL chemiluminescence reagent (EMD Millipore), and images captured on a Bio-Rad ChemiDoc Touch. When necessary, membranes were stripped using 15 g/L glycine, 0.1% SDS, 1% Tween 20, pH 2.2, then re-probed with anti-actin as a control for protein loading. Densitometry was performed using Bio-Rad Image Lab software Version 5.2. A common loading control sample was included on every gel to control for variation in relative band intensities on different blots. Full blot images for each antibody are shown in Supplementary Figure 1.

### Lipid quantification by liquid chromatography-tandem mass spectrometry (LC-MS/MS)

Lipids were extracted from RIPA lysates of the CC (∼100 µg protein) using a two-phase methyl-tert-butyl ether (MTBE):methanol:water (10:3:2.5 v/v/v) extraction procedure (Couttas et al., 2020; Matyash et al., 2008). Internal standards (2 nmole each of d18:1/17:0 ceramide, and d18:1/12:0 sulfatide, d18:1/12:0 HexCer, 1 nmole d18:1/12:0 SM, and 200 pmoles of d17:1 sphingosine and d17:1 S1P) were added at the start of the extraction procedure. Lipids were detected by multiple reaction monitoring on a TSQ Altis mass spectrometer with Vanquish HPLC (ThermoFisher Scientific), operating in positive ion mode. The [M + H]^+^ mass-to-charge (*m/z*) ratio was used for all precursor ions. Product ions were *m/z* 184.1 for SM, 250.2 for d17:1 sphingosine and d17:1 S1P, and 264.3 for all other sphingolipids. Lipids were resolved on a 3 150 mm XDB-C8 column (5 μM particle size) (Agilent) at flow rate 0.4 mL/min. Mobile phase A was 0.2% formic acid, 2 mM ammonium formate in water; and B was 0.2% formic acid, 2 mM ammonium formate in methanol. Total run time was 24 min, starting at 80% B and holding for 2 min, increasing to 100% B from 2 - 14 min, holding at 100% until 20.5 min, then returning to 80% B at 21 min, and holding at 80% B for a further 3 min. Peaks were integrated using TraceFinder 4.1 software (ThermoFisher) and expressed as ratios to the relevant internal standard. Lipid levels were determined with reference to standard curves created by extracting one or two lipid standards from each lipid class, spiked at five different concentrations into 1% (w/v) fatty acid free BSA. Internal standards were added at fixed concentration to the external standards.

### Scanning Electron Microscopy

Male SphK2^-/-^ mice or their WT littermates, at 2 or 15 months of age, were perfused trans- cardially with sterile PBS, then fixative (2.5% glutaraldehyde, 1% paraformaldehyde in 0.15 M PBS). Brains were removed and post-fixed overnight with the same fixative, then stored at 4 °C in 30% sucrose cyroprotectant. The brain was serially sectioned into 1 mm thick coronal sections. Blocks (1 mm x 2 mm) were dissected from the coronal section located 7 mm posterior to the olfactory bulbs. Blocks were immersion fixed for 1 h in 2.5% glutaraldehyde, 2% paraformaldehyde in 0.1 M phosphate buffer, washed in 0.1 M phosphate buffer, then subjected to secondary fixation in 1% osmium tetroxide in 0.1 M phosphate buffer, followed by dehydration through graded alcohols, and Epon (medium) resin infiltration. Following polymerization for 48 h at 60 °C, blocks were sectioned at 90 nm thickness on an ultramicrotome (Ultracut 7, Leica), mounted on copper grids, and post-stained in 2% uranyl acetate and Reynold’s lead citrate. Images were captured using a backscatter detector on a scanning electron microscope (Zeiss Sigma HD FEG SEM). G-ratios were calculated as the axon lumen diameter divided by the outer diameter (axon lumen plus myelin sheath). These measurements were taken from 30-60 axons per region per mouse, with 3 mice per age/genotype, using ImageJ software. Unmyelinated axons were identified by their pale axoplasm containing neurofilaments and microtubules, with an absence of clear myelin sheaths surrounding the axolemma. Measurements were taken perpendicular to the longest axis of axons in cross section, to compensate for axons cross-sectioned in an oblique plane. The operator was blinded to sample groups.

### Statistical Analysis

Results arising from the cuprizone-treated mice were analysed using two-way ANOVAs, with genotype (WT or SphK2^-/-^) and treatment (Ctrl, 6W, 6+2W) as the independent variables. Bonferroni’s post-hoc test was used to assess the effect of different treatments within each genotype, or the effect of genotype under each treatment condition. For the 4 week remyelination experiment using only SphK2^-/-^ mice, one-way ANOVA was used. The Anderson-Darling test was used to assess residual normality. If the residuals were not normally distributed, a natural log transform of the data was performed prior to ANOVA. This resulted in a normal distribution for all analyses except for CNP data in Figure 2B and sphingosine data in Figure 6C. In these cases, analysis of normality in individual treatment groups using the Shapiro-Wilk test indicated normally distributed data prior to ANOVA.

For measurements of mean g-ratio, axon diameter, and proportion of myelinated axons, a two-way ANOVA was used with genotype and age as the independent variables. Least squares regression was used to create lines of best fit for g-ratio against axon diameter (no weighting), and the sum-of-squares F test was applied to determine if the fit of the lines differed significantly between WT and SphK2^-/-^ mice. These analyses were all performed using GraphPad PRISM.

The *car* and *multcomp* packages in R were used to run a two-way ANOVA on each lipid shown in Table 2, and *p* values were adjusted for multiple comparisons with the Benjamini-Hochberg false discovery rate correction.

**Table 2.** Levels of individual sphingolipid species in CC. Results of two-way ANOVA show both unadjusted *p* values, and those adjusted for multiple comparisons (*q* values) using the Benjamini-Hochberg correction. Bonferonni post-test was used to compare WT and SphK2*^-/-^* groups within each treatment group: \**p* < 0.05, ** *p* < 0.01; *** *p* < 0.001. Sph: sphingosine; Cer: ceramide; HexCer, hexosylceramide; SM, sphingomyelin, ST, sulfatide.

|  | Mean ± SEM (pmoles/mg protein) |  |  |  |  |  | Two-way ANOVA |  |  |  |  |  |  |  |  |
| --- | --- | --- | --- | --- | --- | --- | --- | --- | --- | --- | --- | --- | --- | --- | --- |
|  | Ctrl |  | 6W |  | 6+2W |  | Interaction |  |  | Genotype |  |  | Treatment |  |  |
|  | WT | SphK2<sup>-/-</sup> | WT | SphK2<sup>-/-</sup> | WT | SphK2<sup>-/-</sup> | F | p | q | F | p | q | F | p | q |
| Sph 18:1 | 25.2 ± 2 | 38.7 ± 1.1 | 29.2 ± 1.6 | 124 ± 40.2 *** | 25.8 ± 1.6 | 81.2 ± 3.5 * | 3.53 | 4.1E-02 | 2.2E-01 | 20.20 | 7.7E-05 | 2.9E-03 | 4.27 | 2.2E-02 | 2.9E-02 |
| S1P 18:1 | 93.8 ± 4.2 | 13.3 ± 1.4 ** | 67.2 ± 9 | 130.1 ± 34.5 * | 110.1 ± 9.1 | 136.2 ± 10.3 | 12.00 | 1.1E-04 | 4.3E-03 | 0.06 | 8.1E-01 | 8.6E-01 | 11.36 | 1.7E-04 | 3.5E-04 |
| Cer (d18:1/16:0) | 64.4 ± 6.9 | 58.4 ± 6.9 | 132.9 ± 30.1 | 106.9 ± 19.7 | 38.4 ± 3.4 | 55.1 ± 5.2 | 1.00 | 3.8E-01 | 6.0E-01 | 0.16 | 6.9E-01 | 8.0E-01 | 12.56 | 8.2E-05 | 1.8E-04 |
| Cer (d18:1/18:0) | 509.9 ± 64.1 | 753 ± 33.9 | 730.8 ± 75.5 | 928.6 ± 197.9 | 311 ± 21.2 | 795.1 ± 86 ** | 1.48 | 2.4E-01 | 4.4E-01 | 16.74 | 2.5E-04 | 4.0E-03 | 4.91 | 1.3E-02 | 1.9E-02 |
| Cer (d18:1/20:0) | 1.6 ± 0.2 | 2.4 ± 0.7 | 2.9 ± 0.6 | 4.8 ± 0.9 | 2.7 ± 0.6 | 7.5 ± 2 ** | 2.02 | 1.5E-01 | 3.6E-01 | 8.73 | 5.7E-03 | 3.1E-02 | 4.67 | 1.6E-02 | 2.2E-02 |
| Cer (d18:1/22:1) | 8.6 ± 1.2 | 8.3 ± 1.1 | 7 ± 1.6 | 3.5 ± 1 | 2 ± 0.4 | 2 ± 0.6 | 1.79 | 1.8E-01 | 3.9E-01 | 2.21 | 1.5E-01 | 2.3E-01 | 19.87 | 1.9E-06 | 1.0E-05 |
| Cer (d18:1/24:0) | 0.7 ± 0.2 | 0.5 ± 0.1 | 0.7 ± 0.1 | 0.8 ± 0.2 | 0.5 ± 0.1 | 0.9 ± 0.2 | 1.62 | 2.1E-01 | 4.2E-01 | 0.45 | 5.1E-01 | 6.0E-01 | 0.40 | 6.7E-01 | 6.7E-01 |
| Cer (d18:1/24:1) | 58 ± 5.8 | 45 ± 5.9 | 94.4 ± 14.1 | 75.6 ± 8.2 | 31.2 ± 3.2 | 44.5 ± 6.5 | 2.36 | 1.1E-01 | 3.5E-01 | 0.86 | 3.6E-01 | 4.4E-01 | 18.38 | 3.9E-06 | 1.7E-05 |
| HexCer (d18:1/16:0) | 428.5 ± 13.1 | 422.5 ± 37 | 389.4 ± 13.6 | 427.6 ± 29.8 | 401.4 ± 15.8 | 504.2 ± 44.7 * | 1.93 | 1.6E-01 | 3.6E-01 | 3.81 | 5.9E-02 | 1.2E-01 | 1.33 | 2.8E-01 | 3.0E-01 |
| HexCer (d18:1/18:0) | 2652.4 ± 62.3 | 2960.2 ± 160.9 | 2747.7 ± 302.1 | 3289 ± 341.2 | 3503.4 ± 260.4 | 5588.8 ± 436.5 *** | 5.51 | 8.4E-03 | 8.3E-02 | 16.06 | 3.2E-04 | 4.0E-03 | 21.30 | 1.0E-06 | 1.0E-05 |
| HexCer (d18:1/18:0-OH) | 739.3 ± 43 | 994.3 ± 114.2 | 858.3 ± 137.3 | 1037.2 ± 170.8 | 1140.1 ± 142.6 | 2181.4 ± 270.6 *** | 4.30 | 2.2E-02 | 1.5E-01 | 12.84 | 1.0E-03 | 8.0E-03 | 14.30 | 3.1E-05 | 9.1E-05 |
| HexCer (d18:1/20:0) | 382.4 ± 22.7 | 421.8 ± 55.2 | 413.6 ± 66.2 | 428.6 ± 66.4 | 504.8 ± 61.9 | 675.4 ± 74.9 | 0.96 | 3.9E-01 | 6.0E-01 | 2.16 | 1.5E-01 | 2.3E-01 | 5.81 | 6.7E-03 | 1.0E-02 |
| HexCer (d18:1/22:0) | 430.8 ± 48.4 | 587.5 ± 126.7 | 835.4 ± 156 | 993.1 ± 187.3 | 1127.8 ± 174.8 | 1955.2 ± 248.7 ** | 2.57 | 9.1E-02 | 3.5E-01 | 7.02 | 1.2E-02 | 3.9E-02 | 17.98 | 4.7E-06 | 1.7E-05 |
| HexCer (d18:1/22:0-OH) | 464.1 ± 52.1 | 941 ± 295.3 | 978.4 ± 205.8 | 1153.2 ± 291.7 | 1345.6 ± 270.3 | 2150.3 ± 325.5 | 0.75 | 4.8E-01 | 6.4E-01 | 5.09 | 3.1E-02 | 7.8E-02 | 8.39 | 1.1E-03 | 2.0E-03 |
| HexCer (d18:1/22:1) | 2129.8 ± 114 | 2110.4 ± 162.1 | 1461.6 ± 182.6 | 993.8 ± 164.6 | 1570.1 ± 118.9 | 1151.8 ± 123.1 | 1.33 | 2.8E-01 | 4.6E-01 | 6.35 | 1.7E-02 | 4.8E-02 | 20.38 | 1.5E-06 | 1.0E-05 |
| HexCer (d18:1/22:1-OH) | 934.5 ± 52.2 | 1247 ± 108.3 | 721.3 ± 93 | 587.8 ± 107.1 | 797.9 ± 76 | 669.3 ± 70.5 | 4.19 | 2.4E-02 | 1.5E-01 | 0.06 | 8.1E-01 | 8.6E-01 | 13.65 | 4.4E-05 | 1.1E-04 |
| HexCer (d18:1/24:0) | 376.3 ± 48.7 | 559.1 ± 142 | 862.7 ± 139.1 | 1049.8 ± 253 | 1280 ± 236.2 | 2264.5 ± 349.7 ** | 2.17 | 1.3E-01 | 3.5E-01 | 5.87 | 2.1E-02 | 5.7E-02 | 17.20 | 6.9E-06 | 2.2E-05 |
| HexCer (d18:1/24:0-OH) | 287.9 ± 31.7 | 542.2 ± 160.2 | 680.5 ± 135.2 | 643.6 ± 185.2 | 878.2 ± 170.2 | 1207 ± 212.6 | 0.70 | 5.1E-01 | 6.4E-01 | 1.82 | 1.9E-01 | 2.6E-01 | 7.53 | 2.0E-03 | 3.2E-03 |
| HexCer (d18:1/24:1) | 25932.9 ± 1741 | 23620 ± 2359.1 | 25015.6 ± 3145.3 | 18423.7 ± 3269.8 | 30242.1 ± 3075.8 | 20850.5 ± 2145.8 * | 0.83 | 4.5E-01 | 6.3E-01 | 7.28 | 1.1E-02 | 3.7E-02 | 1.09 | 3.5E-01 | 3.7E-01 |
| HexCer (d18:1/24:1-OH) | 3662.4 ± 308.3 | 4591.2 ± 650.4 | 4349.7 ± 687.5 | 3923.4 ± 730.1 | 5219.4 ± 715.9 | 4491.4 ± 534.4 | 0.92 | 4.1E-01 | 6.0E-01 | 0.02 | 8.9E-01 | 8.9E-01 | 0.90 | 4.1E-01 | 4.3E-01 |
| SM (d18:1/16:0) | 2136.4 ± 71.9 | 2022.4 ± 85.4 | 3354 ± 296 | 3828.1 ± 294 | 3341.8 ± 97.5 | 4119.9 ± 333 * | 1.93 | 1.6E-01 | 3.6E-01 | 4.09 | 5.1E-02 | 1.2E-01 | 30.52 | 2.6E-08 | 5.1E-07 |
| SM (d18:1/18:0) | 11535.3 ± 326.7 | 11291.4 ± 415.2 | 12360.6 ± 688 | 12785.5 ± 877.1 | 13110.6 ± 377.3 | 15196.9 ± 1055.2 | 1.58 | 2.2E-01 | 4.2E-01 | 1.81 | 1.9E-01 | 2.6E-01 | 8.19 | 1.3E-03 | 2.2E-03 |
| SM (d18:1/20:0) | 1246.7 ± 84 | 1069.9 ± 65.9 | 1437.7 ± 55.5 | 1424.9 ± 106.3 | 1574.4 ± 48.7 | 1396.9 ± 109.6 | 0.69 | 5.1E-01 | 6.4E-01 | 3.47 | 7.1E-02 | 1.4E-01 | 9.23 | 6.3E-04 | 1.2E-03 |
| SM (d18:1/22:0) | 1644.7 ± 179.3 | 1423.2 ± 123.1 | 2454.1 ± 223.8 | 3095.6 ± 323.7 | 2350.2 ± 109.7 | 2949.6 ± 244.2 | 2.56 | 9.2E-02 | 3.5E-01 | 3.94 | 5.5E-02 | 1.2E-01 | 20.15 | 1.7E-06 | 1.0E-05 |
| SM (d18:1/22:1) | 3273.7 ± 339 | 3823.6 ± 467 | 3155.8 ± 320.4 | 4515.4 ± 696.6 | 2379.7 ± 113 | 3056.3 ± 248.4 | 0.65 | 5.3E-01 | 6.4E-01 | 7.76 | 8.7E-03 | 3.7E-02 | 5.00 | 1.2E-02 | 1.8E-02 |
| SM (d18:1/24:0) | 595 ± 58.5 | 568 ± 52.2 | 990 ± 83.5 | 1369.4 ± 154.2 | 1184.6 ± 76.6 | 1665.9 ± 170 ** | 2.88 | 7.0E-02 | 3.3E-01 | 9.49 | 4.1E-03 | 2.6E-02 | 30.38 | 2.7E-08 | 5.1E-07 |
| SM (d18:1/24:1) | 22954.7 ± 723.5 | 20763.1 ± 1486.2 | 20862.9 ± 1339.7 | 18576 ± 1600.6 | 22772.3 ± 933.9 | 20652.3 ± 1803.5 | 0.00 | 1.0E+00 | 1.0E+00 | 3.87 | 5.7E-02 | 1.2E-01 | 1.50 | 2.4E-01 | 2.6E-01 |
| ST (d18:1/16:0) | 12.3 ± 1 | 9.6 ± 0.8 | 11.6 ± 0.9 | 11.6 ± 0.8 | 13.5 ± 0.8 | 20.7 ± 3.2 ** | 5.46 | 8.8E-03 | 8.3E-02 | 1.29 | 2.6E-01 | 3.3E-01 | 9.48 | 5.4E-04 | 1.1E-03 |
| ST (d18:1/18:0) | 425.7 ± 11.1 | 421.4 ± 28.4 | 354.7 ± 31.9 | 296.3 ± 26.4 | 410.3 ± 24.8 | 572.3 ± 54.1 ** | 6.33 | 4.6E-03 | 8.3E-02 | 1.48 | 2.3E-01 | 3.0E-01 | 13.06 | 6.2E-05 | 1.5E-04 |
| ST (d18:1/20:0) | 108.1 ± 7.1 | 100.1 ± 5.9 | 79.3 ± 12.3 | 59 ± 11.6 | 93 ± 8.7 | 85.7 ± 10.8 | 0.27 | 7.6E-01 | 8.1E-01 | 2.14 | 1.5E-01 | 2.3E-01 | 5.93 | 6.2E-03 | 9.8E-03 |
| ST (d18:1/22:0) | 233.5 ± 13.7 | 225.4 ± 16 | 190.1 ± 36.6 | 157 ± 26 | 233.9 ± 23.4 | 257 ± 35.4 | 0.54 | 5.9E-01 | 6.7E-01 | 0.07 | 7.9E-01 | 8.6E-01 | 3.76 | 3.3E-02 | 4.1E-02 |
| ST (d18:1/22:0-OH) | 56.1 ± 4.5 | 78.5 ± 13 | 57.4 ± 11.5 | 48.4 ± 7.2 | 67 ± 9.2 | 85.3 ± 10.8 | 1.42 | 2.5E-01 | 4.4E-01 | 1.68 | 2.0E-01 | 2.8E-01 | 2.87 | 7.0E-02 | 8.4E-02 |
| ST (d18:1/22:1) | 243.4 ± 11 | 214.3 ± 14.4 | 150.1 ± 21.1 | 108.6 ± 21.4 | 185.3 ± 12.1 | 139.8 ± 15.1 | 0.13 | 8.8E-01 | 9.0E-01 | 8.36 | 6.7E-03 | 3.2E-02 | 18.01 | 4.6E-06 | 1.7E-05 |
| ST (d18:1/22:1-OH) | 99.2 ± 6.3 | 107.4 ± 5.6 | 63.8 ± 8.1 | 51.9 ± 9.1 | 85.4 ± 7.9 | 60.7 ± 6.4 | 2.44 | 1.0E-01 | 3.5E-01 | 2.40 | 1.3E-01 | 2.3E-01 | 17.91 | 4.9E-06 | 1.7E-05 |
| ST (d18:1/24:0) | 308.3 ± 21.9 | 331.8 ± 34.3 | 325.7 ± 59.7 | 282.4 ± 49.9 | 412 ± 47.3 | 449.3 ± 59.1 | 0.39 | 6.8E-01 | 7.4E-01 | 0.02 | 8.9E-01 | 8.9E-01 | 4.16 | 2.4E-02 | 3.1E-02 |
| ST (d18:1/24:0-OH) | 34.1 ± 4.2 | 48.4 ± 11.5 | 39.7 ± 7.8 | 39.4 ± 8.6 | 47.2 ± 7.5 | 65.4 ± 10.6 | 0.63 | 5.4E-01 | 6.4E-01 | 2.28 | 1.4E-01 | 2.3E-01 | 2.36 | 1.1E-01 | 1.3E-01 |
| ST (d18:1/24:1) | 4793.5 ± 187.8 | 4088.5 ± 288.7 | 3142.6 ± 427.1 | 2194.4 ± 356.5 | 4042.1 ± 300.2 | 2707 ± 307.1 * | 0.48 | 6.2E-01 | 6.9E-01 | 13.87 | 7.1E-04 | 6.7E-03 | 13.97 | 3.7E-05 | 1.0E-04 |
| ST (d18:1/24:1-OH) | 380.8 ± 15.9 | 381.9 ± 29.7 | 252.4 ± 35.1 | 182.7 ± 28.1 | 297.2 ± 25.1 | 190 ± 16.2 * | 2.16 | 1.3E-01 | 3.5E-01 | 7.42 | 1.0E-02 | 3.7E-02 | 21.09 | 1.1E-06 | 1.0E-05 |

### Data Availability

The data that support the findings of this study are available from the corresponding author upon reasonable request.

## Results

### SphK2 is essential for remyelination following cuprizone withdrawal

To determine if SphK2 is required for protection of oligodendrocytes and myelin, and for spontaneous remyelination following cuprizone withdrawal, 12 week old SphK2^-/-^ mice and their WT littermates were fed cuprizone for 6 weeks and culled immediately (6W), or returned to a normal chow diet (i.e. removal of cuprizone from the diet) for 2 weeks prior to culling (6+2W). These were compared to mice fed a normal chow diet for 6 weeks (Ctrl). Luxol fast blue staining showed robust demyelination in the corpus callosum (CC) of 6W WT compared to Ctrl WT mice, and remyelination in 6+2W WT mice (Figure 1a). Demyelination was more pronounced in 6W SphK2^-/-^ compared to 6W WT mice, and remyelination was not apparent in 6+2W SphK2^-/-^ mice (Figure 1a).

**Figure 1.**
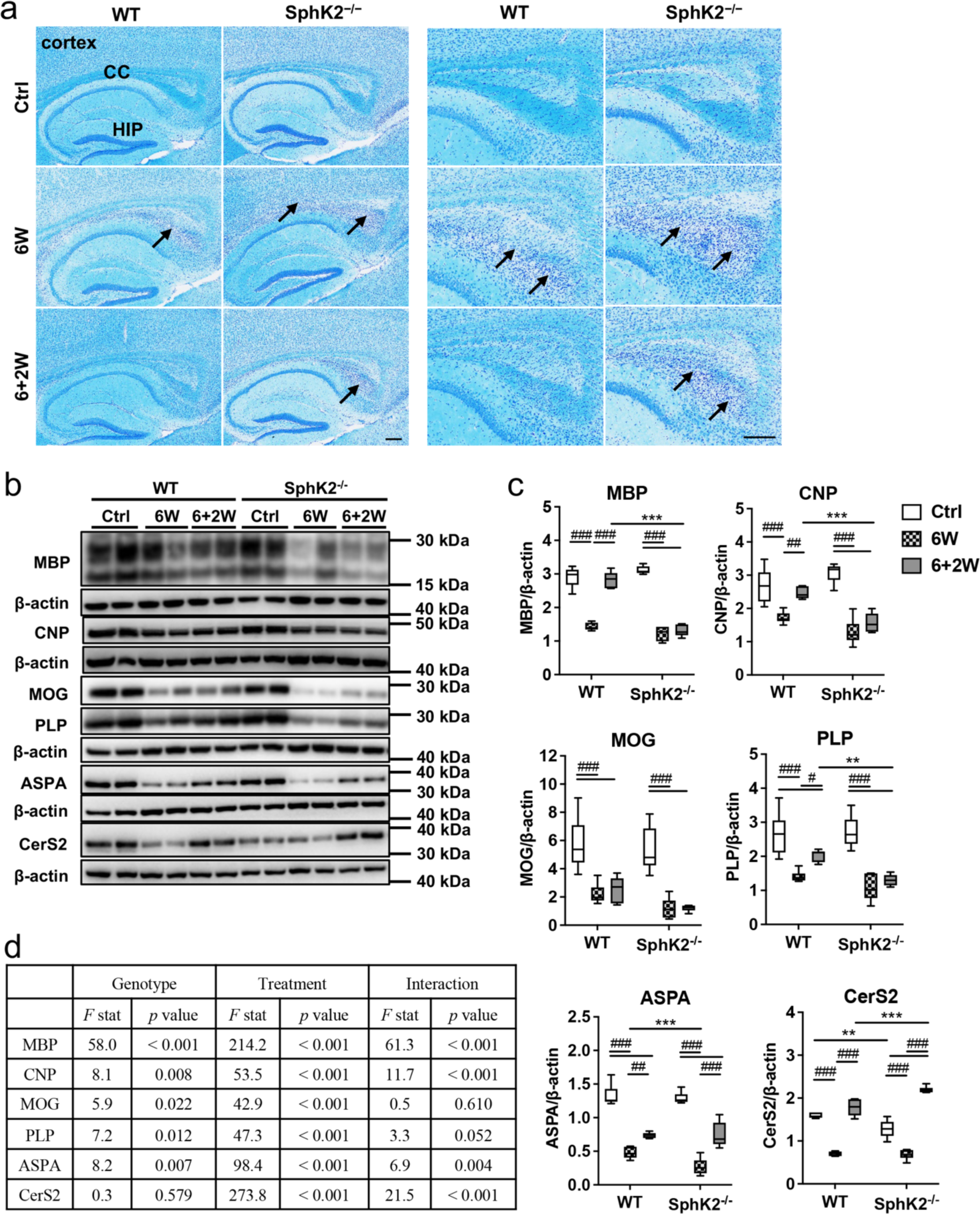
SphK2 deletion impairs remyelination following cuprizone withdrawal. (a) Luxol fast blue (light blue)/cresyl violet (purple) staining of sagittal sections from WT or SphK2^−/−^ mice fed control chow (Ctrl) or cuprizone for 6 weeks (6W), or cuprizone for 6 weeks then normal chow for 2 weeks (6+2W). Images on left show the body and splenium of the CC above the hippocampus (HIP). Images on right show an enlarged view of the splenium, where demyelination is clearly apparent (marked by arrows). Scale bar, 200 μm. (b) Western blots, (c) densitometric quantification, and (d) results of two-way ANOVA for myelin and oligodendrocyte markers in the CC. Protein levels were normalised to β-actin in each sample, n = 6 mice/group. Box and whisker plots show full data range with 25^th^ – 75^th^ percentile boxed, and horizontal bar marking the median. Ctrl, open boxes; 6W, hatched boxes; 6+2W, grey boxes. Statistical significance was determined by two-way ANOVA with results of Bonferroni’s post-tests shown on the graphs: between-genotype comparisons (\**p* < 0.05, ** *p* < 0.01; *** *p* < 0.001) and within-genotype comparisons (# *p* < 0.05, ## *p* < 0.01; ### *p* < 0.001).

To quantify myelin content, we performed western blotting for the myelin markers myelin basic protein (MBP), myelin proteolipid protein (PLP), myelin oligodendrocyte glycoprotein (MOG), and 2’,3’-cyclic nucleotide 3’-phosphodiesterase (CNP) in CC (Figure 1b-d) and cortex (Figure 2a-c) homogenates. Myelin protein levels in the CC did not differ between Ctrl WT and Ctrl SphK2^−/−^ mice at 18 weeks of age, however PLP and MOG levels were significantly lower in the cortex of SphK2^−/−^ mice. Compared to the Ctrl groups, a pronounced loss of all four myelin markers was observed in both CC and cortex of the 6W WT and SphK2^-/-^ groups, and a significant effect of both genotype and treatment was observed for all four markers in two-way ANOVA (Figure 1d and 2c). MBP, CNP and PLP levels were significantly higher in the CC of 6+2W compared to 6W WT mice (*p* < 0.001, *p* = 0.002, and *p* = 0.046, respectively), whereas none of these markers were higher in 6+2W compared to 6W SphK2^-/-^ mice, indicating that remyelination was impaired in SphK2^-/-^ mice. In contrast to the other three myelin markers, MOG levels were not higher in the CC of 6+2W compared to 6W WT mice. This is in agreement with previous studies showing that MOG recovers more slowly than other myelin markers during remyelination in the cuprizone model (Gudi et al., 2009; Lindner et al., 2008).

**Figure 2.**
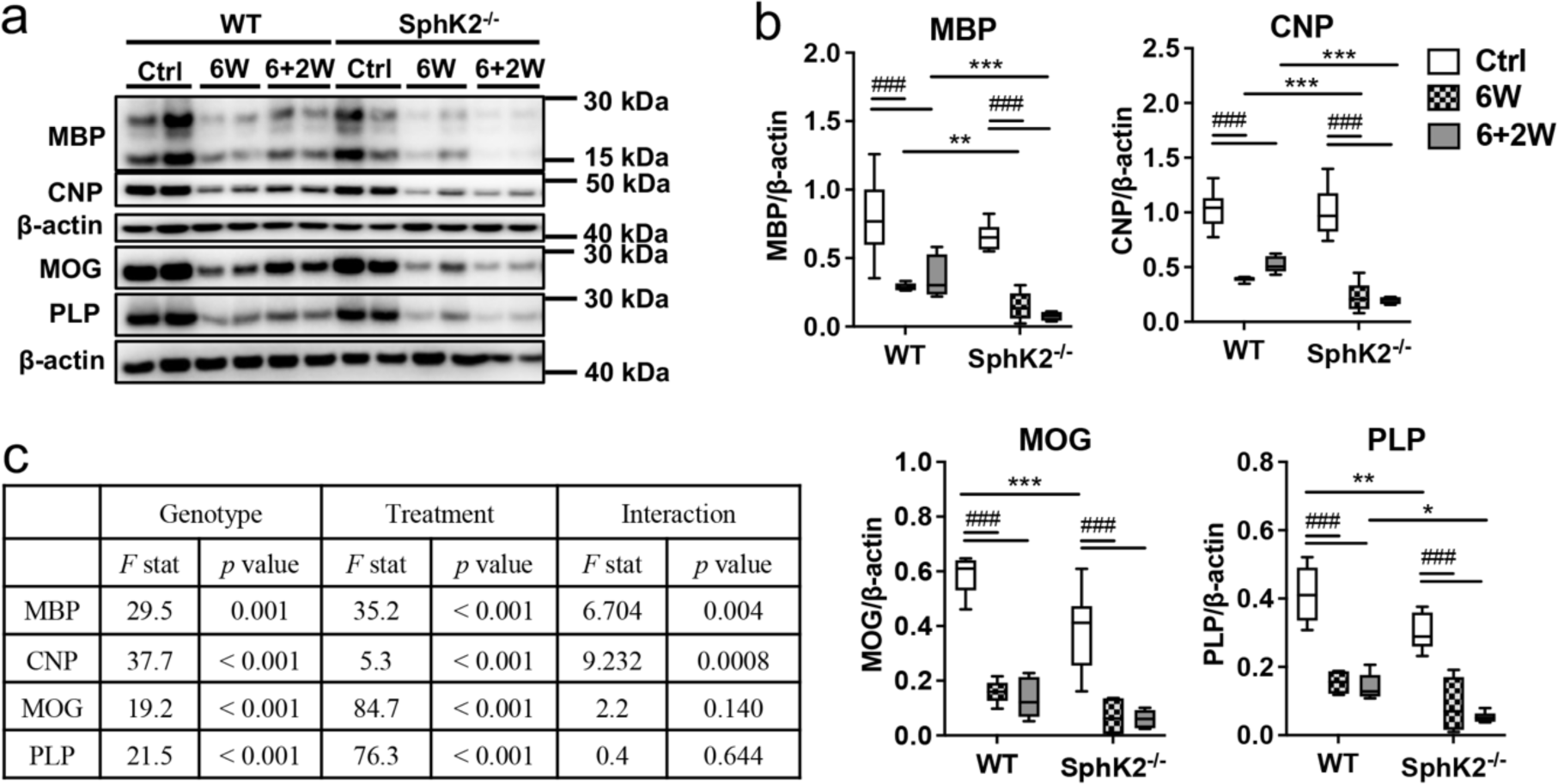
SphK2 protects against myelin loss in the cortex. (a) Western blots, (b) densitometric quantification, and (c) results of two-way ANOVA for myelin and oligodendrocyte markers in the cortex. Protein levels were normalised to β-actin in each sample, n = 5 - 6 mice/group. Box and whisker plots show full data range with 25^th^ – 75^th^ percentile boxed, and horizontal bar marking the median. Results of Bonferroni’s post-tests are shown on the graphs: between-genotype comparisons (\**p* < 0.05, ** *p* < 0.01; *** *p* < 0.001) and within-genotype comparisons (# *p* < 0.05, ## *p* < 0.01; ### *p* < 0.001).

Myelin marker levels were not significantly higher in the cortex of 6+2W compared to 6W groups of either genotype, consistent with previous studies reporting that remyelination after cuprizone withdrawal occurs more slowly in the cortex than the CC (Baxi et al., 2017; Gudi et al., 2009). However, MBP and CNP levels were significantly lower in the cortex of 6W SphK2^- /-^ compared to 6W WT mice, and 6+2W SphK2^-/-^ compared to 6+2W WT mice, indicative of more severe cuprizone-mediated demyelination in the cortex of SphK2^-/-^ mice.

### SphK2 protects against loss of mature oligodendrocytes

We next determined if cuprizone administration differentially affected oligodendrocyte number in WT versus SphK2^-/-^ mice, by counting cells positive for the mature oligodendrocyte marker aspartoacylase (Aspa) (Madhavarao et al., 2004; Y. Zhang et al., 2014) (Figure 3a). Oligodendrocyte density did not differ between Ctrl WT and Ctrl SphK2^-/-^ mice in the CC (Figure 3b) or cortex (Figure 3c). Cuprizone feeding produced a pronounced depletion of Aspa-positive cells in the CC (Figure 3b) and cortex (Figure 3c), which was more pronounced in SphK2^-/-^ compared to WT mice. Aspa-positive cell density was reduced 61% in CC of 6W WT mice, and 84% in 6W SphK2^-/-^, relative to the respective Ctrl groups (2-way ANOVA, effect of genotype: F = 8.2, *p* = 0.008; treatment: F = 87.7, *p* < 0.001; interaction: F = 1.7, *p* = 0.19). In the cortex, Aspa-positive cell density declined 50% in 6W WT and 72% in 6W SphK2^-/-^ mice, relative to the Ctrl groups (2-way ANOVA, effect of genotype: F = 26.7, *p* < 0.001; treatment: F = 56.1, *p* < 0.001; interaction: F = 1.5, *p* = 0.25).

**Figure 3.**
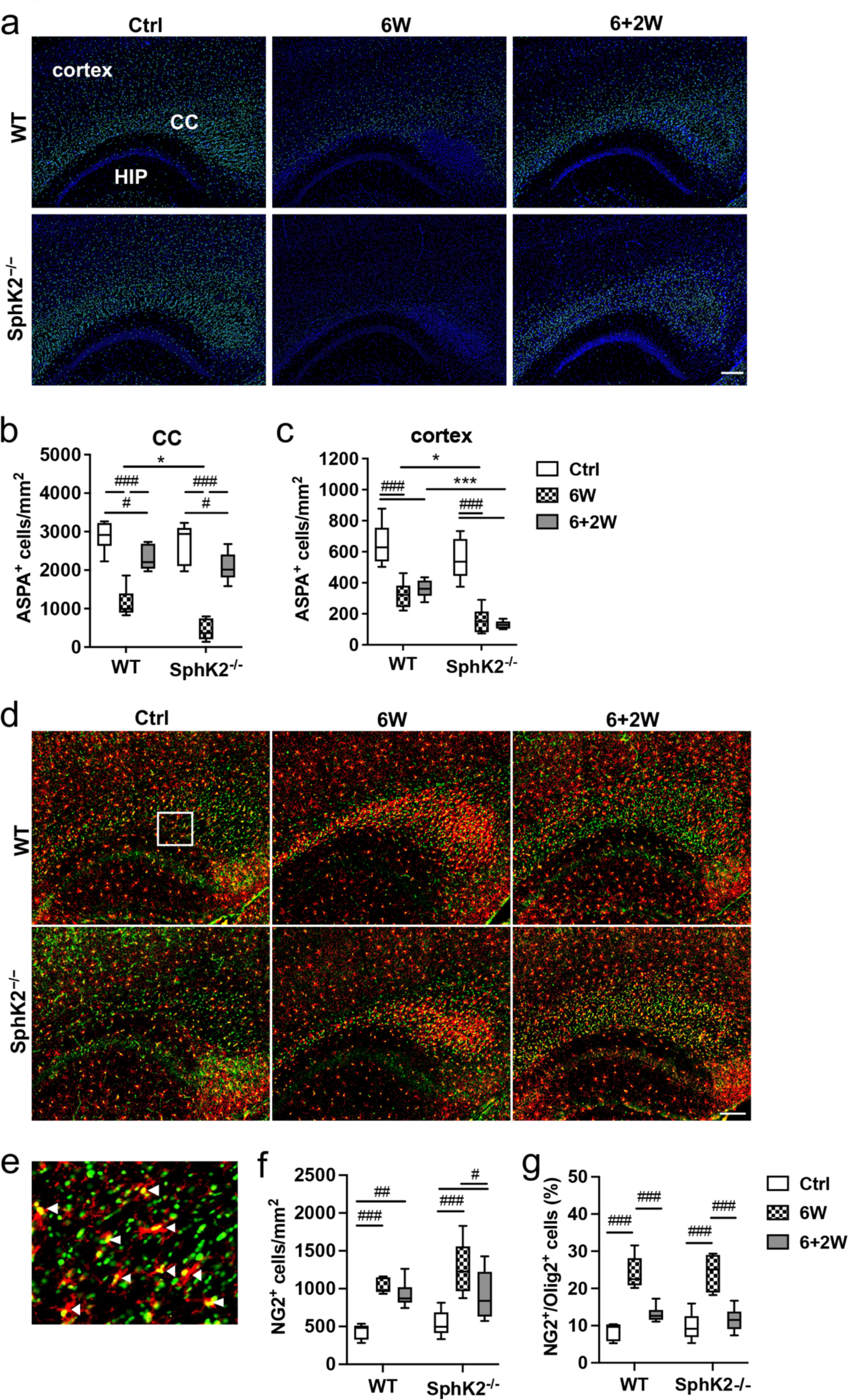
Loss of SphK2 sensitizes to oligodendrocyte loss induced by cuprizone. (a) ASPA (green) and DAPI (blue) immunofluorescent staining, and (b,c) quantified ASPA- positive cell density (cells per mm^2^) in the CC (b) and cortex (c) of WT or SphK2^−/−^ mice (6 mice/group), in Ctrl (clear), 6W (hatched), or 6+2W (grey) treatment groups. Scale bar, 300 μm. (d) Representative immunofluorescence images for Olig2 (green) and NG2 (red). Scale bar, 100 μm. (e) Higher magnification view of the boxed area in (d), illustrating NG2-positive OPCs (arrowheads). (f) Density of cells positive for both NG2 and Olig2 in the CC. (g) Proportion of Olig2-positive cells in the CC that are also positive for NG2. Box and whisker plots show full data range with 25^th^ – 75^th^ percentile boxed, and horizontal bar marking the median (6 mice/group). Statistical significance was determined by two-way ANOVA with results of Bonferroni’s post-tests shown on the graphs: between-genotype comparisons (\**p* < 0.05, *** *p* < 0.001) and within-genotype comparisons (# *p* < 0.05, ## *p* < 0.01; ### *p* < 0.001).

Mature oligodendrocyte density in the CC rebounded following cuprizone withdrawal (*p* < 0.001 for both genotypes comparing 6W to 6+2W) (Figure 3b), reaching 80% of the Ctrl group in WT mice, and 77% of the Ctrl in SphK2^-/-^ mice. In contrast, oligodendrocyte density in the cortex did not recover after 2 weeks of cuprizone withdrawal and remained significantly lower in SphK2^-/-^ compared to WT mice (Figure 3c).

Cuprizone administration is associated with an expansion of oligodendrocyte progenitor cells (OPCs), as a response to loss of mature oligodendrocytes (Gudi et al., 2009; Mason et al., 2000). OPCs were quantified as cells positive for both the pan-oligodendrocyte lineage transcription factor Olig2 and the OPC marker NG2 proteoglycan (Zhan et al., 2020) (Figure 3d,e). Cells positive for both NG2 and Olig2 doubled in 6W WT compared to Ctrl WT mice, and remained elevated in the 6+2W group (Figure 3f). This response was not notably affected by SphK2 deficiency (2-way ANOVA, genotype: F = 2.2, *p* = 0.15; treatment: F = 26.4, *p* < 0.001; interaction: F = 0.93, *p* = 0.41), although the number of NG2/Olig2 double-positive cells declined in the 6+2W relative to the 6W SphK2^-/-^ group (Figure 3f). In both genotypes, the proportion of Olig2-positive cells that are also NG2-positive increased substantially in the 6W group (2-way ANOVA, genotype: F = 0.001, *p* = 0.97; treatment: F = 58.9, *p* < 0.001; interaction: F = 0.43, *p* = 0.66) (Figure 3g). This measure returned to levels that were not significantly different to the untreated controls in the 6+2W groups, presumably resulting from the restoration of mature OLGs.

In agreement with Aspa-positive cell counts, Aspa protein levels in the CC were decreased in the 6W group and partially restored in the 6+2W group, for both WT and SphK2^-/-^ mice (*p* < 0.001 for both WT and SphK2^-/-^ mice, 6W vs 6+2W) (Figure 1b-c). ASPA protein levels were also significantly lower in 6W SphK2-/- compared to 6W WT mice (*p* < 0.001). Another mature oligodendrocyte marker, ceramide synthase 2 (CerS2) (Couttas et al., 2016; Kremser et al., 2013), which is required for myelin lipid synthesis, showed a similar pattern of reduced levels in the 6W group, increasing substantially in the 6+2W group. Overall, these results support the hypothesis that production of new mature oligodendrocytes from OPCs is not affected by SphK2 deficiency, despite the lack of remyelination.

### Absence of remyelination in the CC of SphK2^-/-^ mice four weeks after cuprizone withdrawa

Since mature oligodendrocyte numbers recovered in the CC of SphK2^−/−^ mice after cuprizone withdrawal, we tested if remyelination in the CC of SphK2^−/−^ mice is simply delayed relative to WT mice. Twelve week old SphK2^−/−^ mice were fed chow containing cuprizone for 6 weeks, then allowed 4 weeks for recovery on normal chow before assessing myelin content (6+4W). Western blotting showed a significant decrease of MBP, CNP, MOG and PLP in the CC of the 6W group. However, these markers did not recover in the 6+4W group, whereas levels of the mature oligodendrocyte markers Aspa and CerS2 rebounded significantly (Figure 4a-b). Luxol fast blue staining (Figure 4c,e) and MBP immunohistochemistry (Figure 4d,f) further verified the lack of remyelination in the 6+4W SphK2^−/−^ group. These results suggest that SphK2^−/−^ is essential for remyelination, rather than simply delaying remyelination following cuprizone withdrawal.

**Figure 4.**
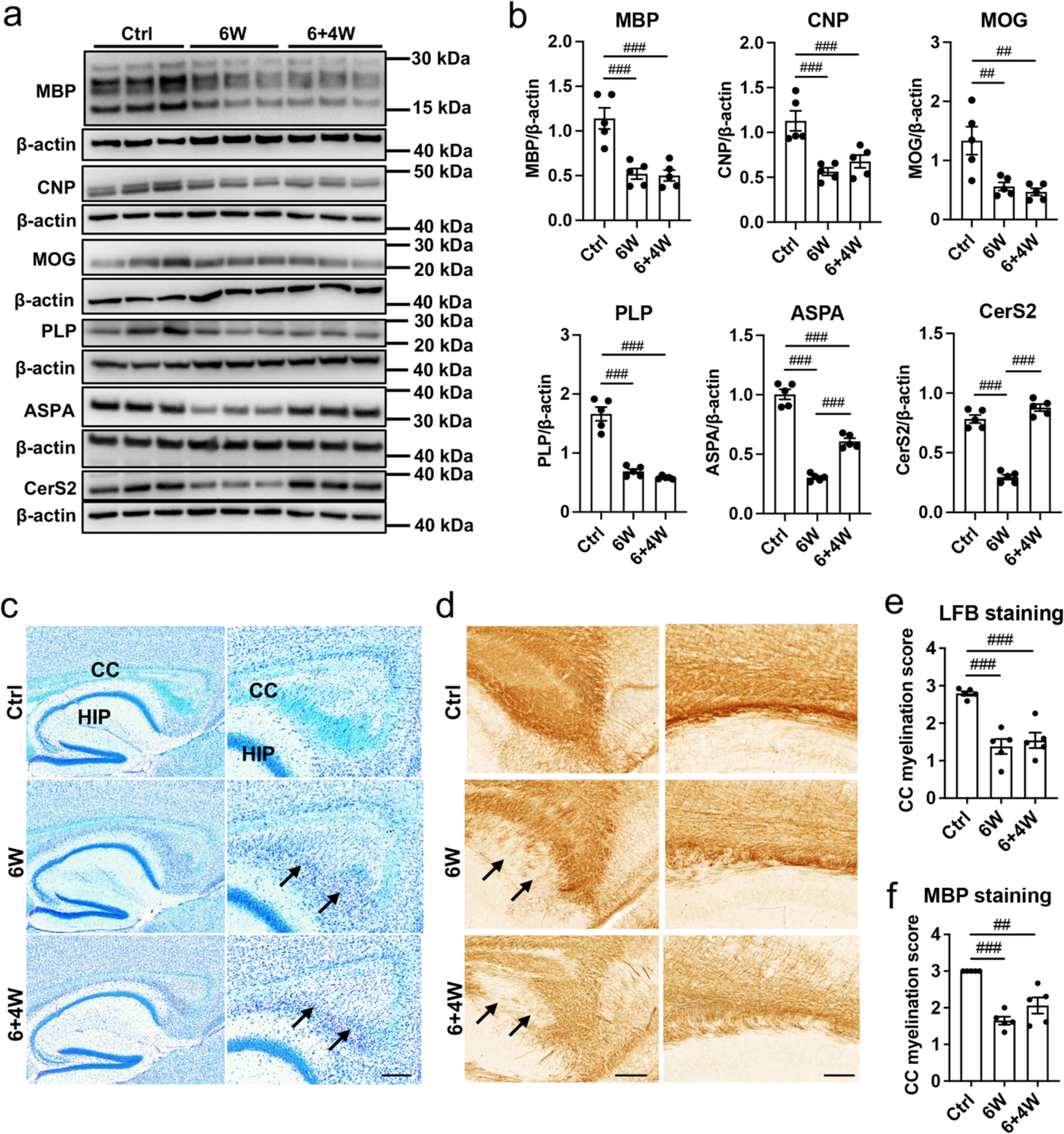
Absence of remyelination 4 weeks after cuprizone withdrawal in SphK2*^−/−^* mice. (a) Western blots and (b) densitometric quantification of myelin and oligodendrocyte markers in the CC of SphK2^-/-^ mice fed normal chow (Ctrl), cuprizone for 6 weeks (6W), or cuprizone for 6 weeks then normal chow for 4 weeks (6+4W). Protein levels were normalised to β-actin in each sample. (c) Representative images for luxol fast blue (LFB)/cresyl violet staining of the CC. HIP: hippocampus. Images on the right are an enlarged view of those on the left, showing the splenium of the CC. (d) Representative images showing MBP staining in the splenium (left) and body (right) of the CC. Demyelination in (c) and (d) is marked by arrows. Scale bar, 200 μm. (e-f) Scoring of CC myelination in (e) LFB and (f) MBP images: 3 = fully myelinated, 0 = fully demyelinated. All graphs show mean ± SEM (n = 5). Statistical significance was determined by one-way ANOVA with Bonferonni’s post-test (## *p* < 0.01; ### *p* < 0.001).

### Astrogliosis does not resolve after cuprizone withdrawal in SphK2^-/-^ mice

Astrogliosis and microgliosis in the CC, visualised with the markers GFAP and Iba1, respectively, are well-established responses to cuprizone feeding (S. Kim et al., 2018; Zhan et al., 2020). A pronounced increase in GFAP (Figure 5a-b) and Iba1 (Figure 5c-d) staining was observed in the CC of cuprizone-fed mice (6W WT and 6W SphK2^-/-^ groups). Relative to the 6W group, GFAP staining subsided in the WT 6+2W group (*p* = 0.0025) but not in the SphK2^- /-^ 6+2W group (*p* > 0.99), indicating deficient resolution of astrogliosis in SphK2^-/-^ mice. Iba1 staining declined significantly in both WT and SphK2^-/-^ mice after cuprizone withdrawal (both genotypes *p* < 0.001, 6W vs 6+2W). Similarly, Iba1 but not GFAP staining was reduced following 4 weeks of cuprizone withdrawal in SphK2^-/-^ mice (i.e. comparing 6+4W to 6W groups) (Figure 5e-g).

**Figure 5.**
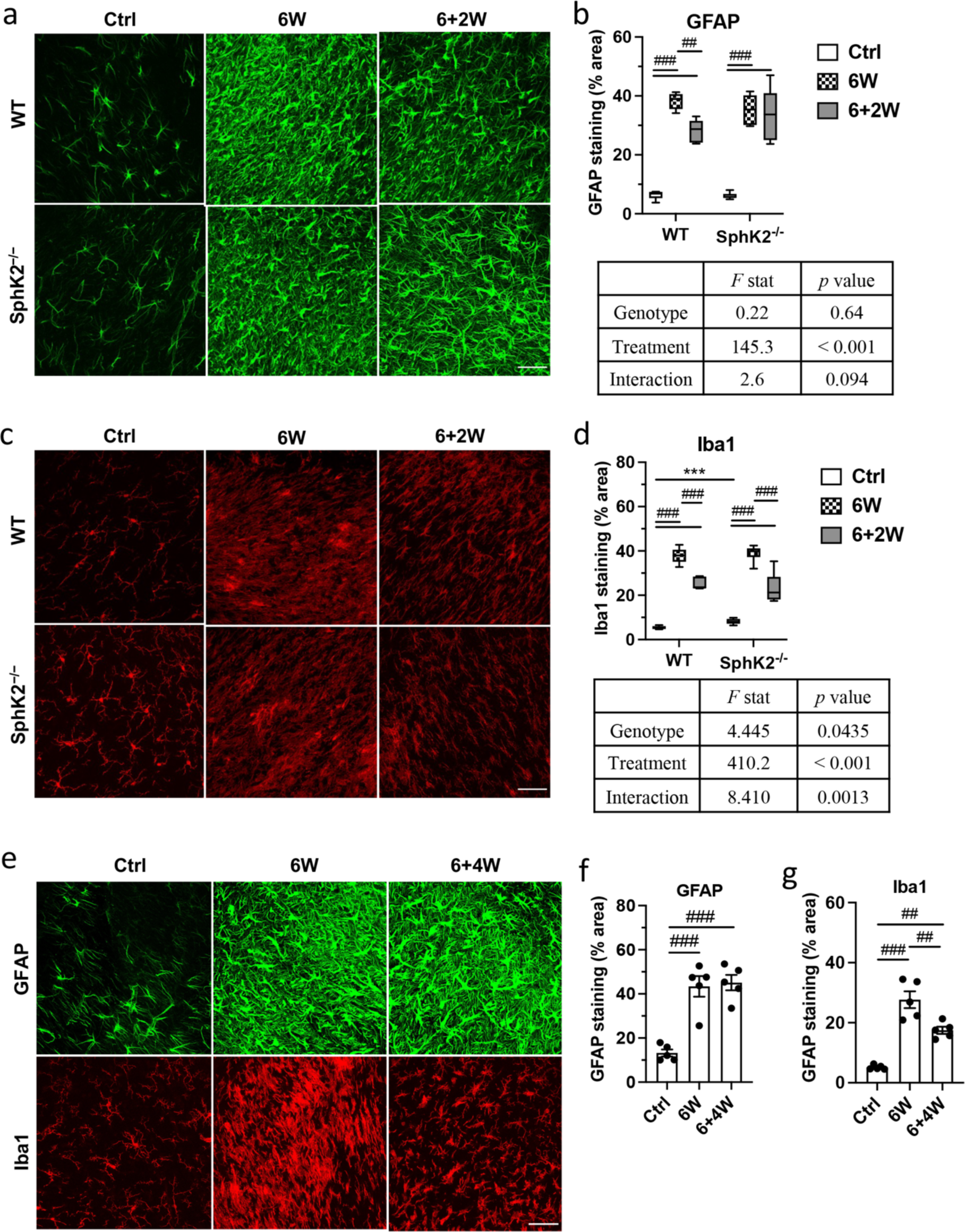
Sustained astrogliosis in SphK2*^-/-^* mice following cuprizone withdrawal. Representative images (a,c) and quantified area of staining (b,d) for GFAP (a,b) and Iba1 (c,d) in the CC of WT or SphK2^-/-^ mice. Scale bar, 50 μm. Box and whisker plots show full data range with 25^th^ – 75^th^ percentile boxed, and horizontal bar marking the median (6 mice/group). Main effect two-way ANOVA results are shown below each graph, and results of Bonferroni’s post-tests are shown on the graphs: between-genotype comparisons (*** *p* < 0.001) and within-genotype comparisons (## *p* < 0.01; ### *p* < 0.001). All comparisons between WT and SphK2^- /-^ mice were non-significant. (e) Representative images and (f,g) quantified area of staining for GFAP (f) and Iba1 (g) in the CC of SphK2^-/-^ mice (5 mice/group). One-way ANOVA with Bonferonni’s post-test was used to assess statistical significance.

### Sphingosine and ceramide levels remain elevated in SphK2^-/-^ mice after cuprizone withdrawal

To determine how the absence of SphK2 affects sphingolipid metabolism during and after cuprizone withdrawal, we quantified levels of S1P, its precursors sphingosine and ceramide, and essential myelin sphingolipids derived from ceramide: sulfatide, hexosylceramide (HexCer), and sphingomyelin (SM) (Figure 6a). Mean S1P levels did not change significantly with cuprizone treatment in WT mice (Figure 6b). S1P levels were 29% lower in the CC of 6W compared to Ctrl WT mice (not significant), and 60% higher in the 6+2W compared to the 6W group (*p* = 0.024). Compared to WT mice, S1P levels were an order of magnitude lower in CC of Ctrl SphK2^-/-^ mice (*p* < 0.001). Remarkably, however, S1P levels were 10-fold higher in the 6W SphK2^-/-^ compared to the Ctrl SphK2^-/-^ group (*p* < 0.001), and remained elevated in the 6+2W group. Sphingosine was unchanged in WT mice across all treatment groups, but increased 3.2-fold in SphK2^-/-^ mice with cuprizone treatment (*p* = 0.003, Ctrl vs 6W). Compared to WT mice, sphingosine content was 4-fold higher in SphK2^-/-^ mice in the 6W treatment group and 3-fold higher in the 6+2W group (both *p* < 0.001) (Figure 6c). Mean ceramide levels were 50% higher in 6W WT compared to Ctrl mice (not significant), and 2.5- fold lower in the 6+2W compared to the 6W group (*p* < 0.001) (Figure 6d). Compared to WT mice, total ceramide levels were not significantly higher in Ctrl or 6W SphK2^-/-^ mice. However, ceramide levels did not decrease in SphK2^-/-^ mice following cuprizone withdrawal, resulting in 2.3-fold higher levels in the CC of 6+2W SphK2^-/-^ compared to 6+2W WT mice (*p* < 0.001).

**Figure 6.**
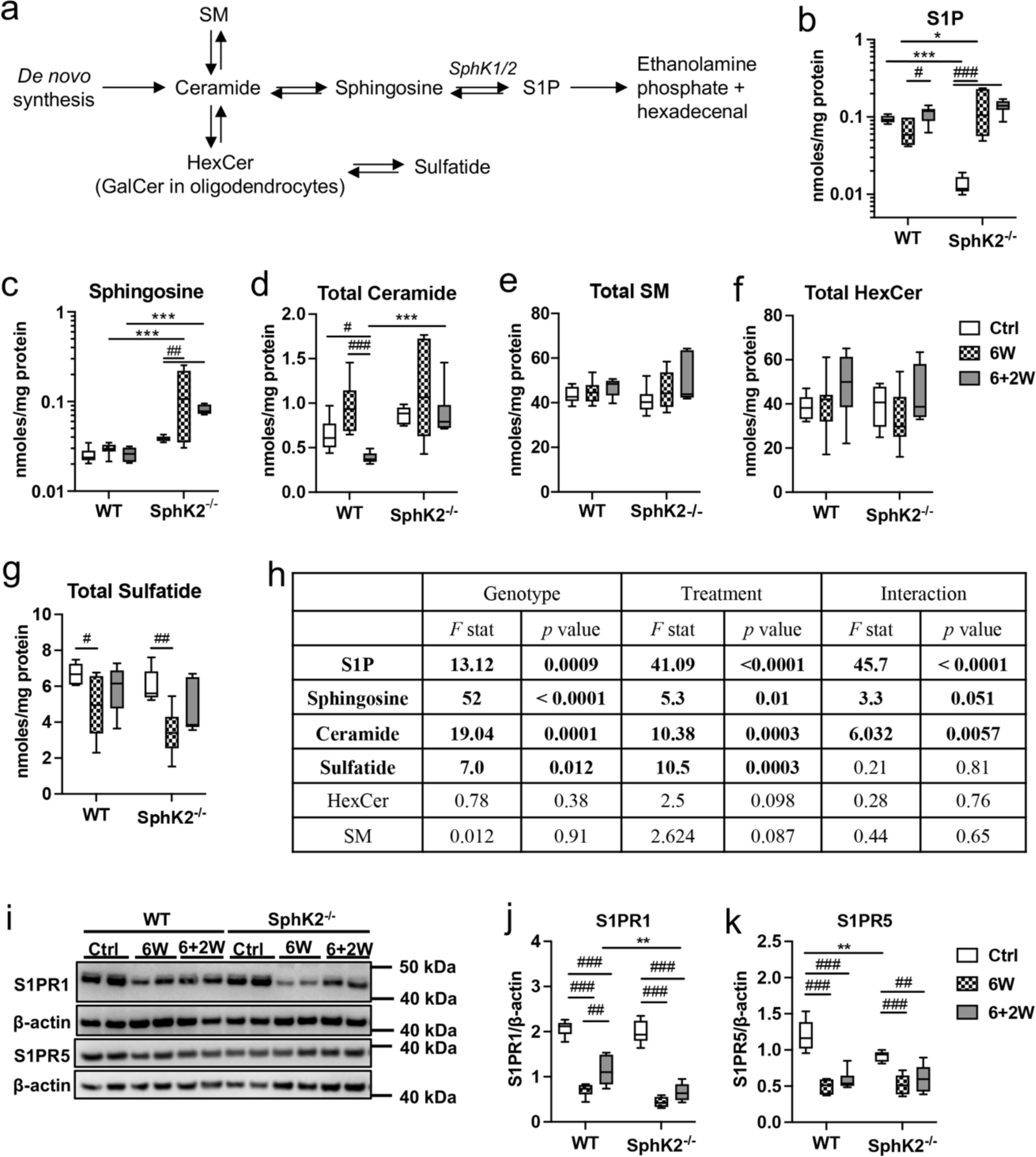
High levels of sphingosine and ceramide in SphK2*^-/-^* mice following cuprizone withdrawal. (a) Basic sphingolipid pathway in oligodendrocytes. SM, sphingomyelin; HexCer, hexosylceramide. HexCer is comprised of both glucosylceramide and galactosylceramide (GalCer). GalCer is an abundant and essential constituent of myelin. (b-g) Levels of (b) S1P, (c) sphingosine, (d) ceramide, (e) SM, (f) HexCer, and (g) sulfatide in CC of WT or SphK2^-/-^ mice, in Ctrl (clear), 6W (hatched), or 6+2W (grey) treatment groups. Box and whisker plots show full data range with 25^th^ – 75^th^ percentile boxed, and horizontal bar marking the median, n = 6 - 8 mice/group. (h) Results of two-way ANOVA for each lipid class. Results of Bonferroni’s post-tests are shown on the graphs: between genotype comparisons (\**p* < 0.05, ** *p* < 0.01; *** *p* < 0.001) and within-genotype comparisons (# *p* < 0.05, ## *p* < 0.01; ### *p* < 0.001). (i) Representative western blots and (j,k) quantified densitometry for (j) S1PR1 and (k) S1PR5 protein levels.

SM and HexCer levels were not significantly affected by treatment or genotype (Figure 6e,f,h), whereas the myelin lipid sulfatide decreased with cuprizone feeding in both WT (30% decrease, *p* = 0.021, 6W compared to Ctrl) and SphK2^-/-^ mice (45% decrease, *p* = 0.003) (Figure 6g). An overall effect of genotype on sulfatide levels was observed (Figure 6h), as levels were lower in SphK2^-/-^ mice. However this difference was not significant in individual treatment group comparisons. Results for individual lipid species are shown in Table 2.

Hyper-activation of S1P receptors, particularly S1PR1, by pharmacological agonists results in their proteosomal degradation and sustained loss of expression (Bigaud et al., 2014; Gonzalez-Cabrera et al., 2007). Western blotting was carried out to determine if the abnormally high S1P levels in cuprizone-treated SphK2^-/-^ mice were associated with decreased levels of S1PR1 or S1PR5 (Figure 6i-k). S1PR1 was significantly regulated by both genotype and treatment (2- way ANOVA, genotype: F = 15.4, *p* < 0.001; treatment: F = 155.8, *p* < 0.001; interaction: F = 2.5, *p* = 0.10), whereas an interaction between genotype and treatment was observed for S1PR5 (genotype: F = 2.5, *p* = 0.13; treatment: F = 46.2, *p* < 0.001; interaction: F = 4.2, *p* = 0.025). Both receptors were strongly down-regulated in the 6W compared to the Ctrl groups. S1PR1 expression increased following cuprizone withdrawal in WT (*p* = 0.004 comparing 6W to 6+2W) but not SphK2^-/-^ mice, and was significantly lower in the 6+2W SphK2^-/-^ compared to the 6+2W WT group (*p* = 0.002). In contrast, S1PR5 was significantly lower in Ctrl SphK2-/- compared to Ctrl WT mice (*p* = 0.008).

### Myelin is thinner in the CC of aged SphK2^-/-^ mice

Since SphK2^-/-^ mice did not remyelinate following cuprizone withdrawal, we hypothesized that these mice may show age-dependent myelin loss, due to impaired physiological myelin turnover. Mature oligodendrocyte density in the CC did not differ between WT and SphK2^-/-^ mice at 20 months of age (Figure 7a). However, analysis of scanning electron microscopy images revealed a significant increase in the g-ratio [diameter of the axon/diameter of (axon + myelin)] in the CC of 15 month old SphK2^-/-^ mice (Figure 7b-e) (2-way ANOVA for mean g-ratio: effect of genotype: F = 34.9, *p* < 0.001; age: F = 61.2, *p* < 0.001; interaction: F = 6.0, *p* = 0.039). The g-ratio did not differ between WT and SphK2^-/-^ mice at 2 months of age. There was no difference in mean axon diameter between WT and SphK2^-/-^ mice at 2 or 15 months of age (Figure 7f) (2-way ANOVA: effect of genotype: F = 0.01, *p* = 0.94; age: F = 4.9, *p* = 0.058; interaction: F = 0.69, *p* = 0.43), confirming that the increased g-ratio in 15 month old SphK2^-/-^ mice is attributed to reduced myelin thickness rather than increased axon diameter. In agreement with this, the line of best fit for g-ratio against axon diameter differed significantly between WT and SphK2^-/-^ mice at 15 months (Figure 7b, *p* < 0.001), but not 2 months of age (Figure 7c). There was no significant difference in the proportion of unmyelinated axons in the CC of WT compared to SphK2^-/-^ mice (Figure 7g) (2-way ANOVA: effect of genotype: F = 5.2, *p* = 0.052; age: F = 0.49, *p* = 0.50; interaction: F = 0.21, *p* = 0.66).

**Figure 7.**
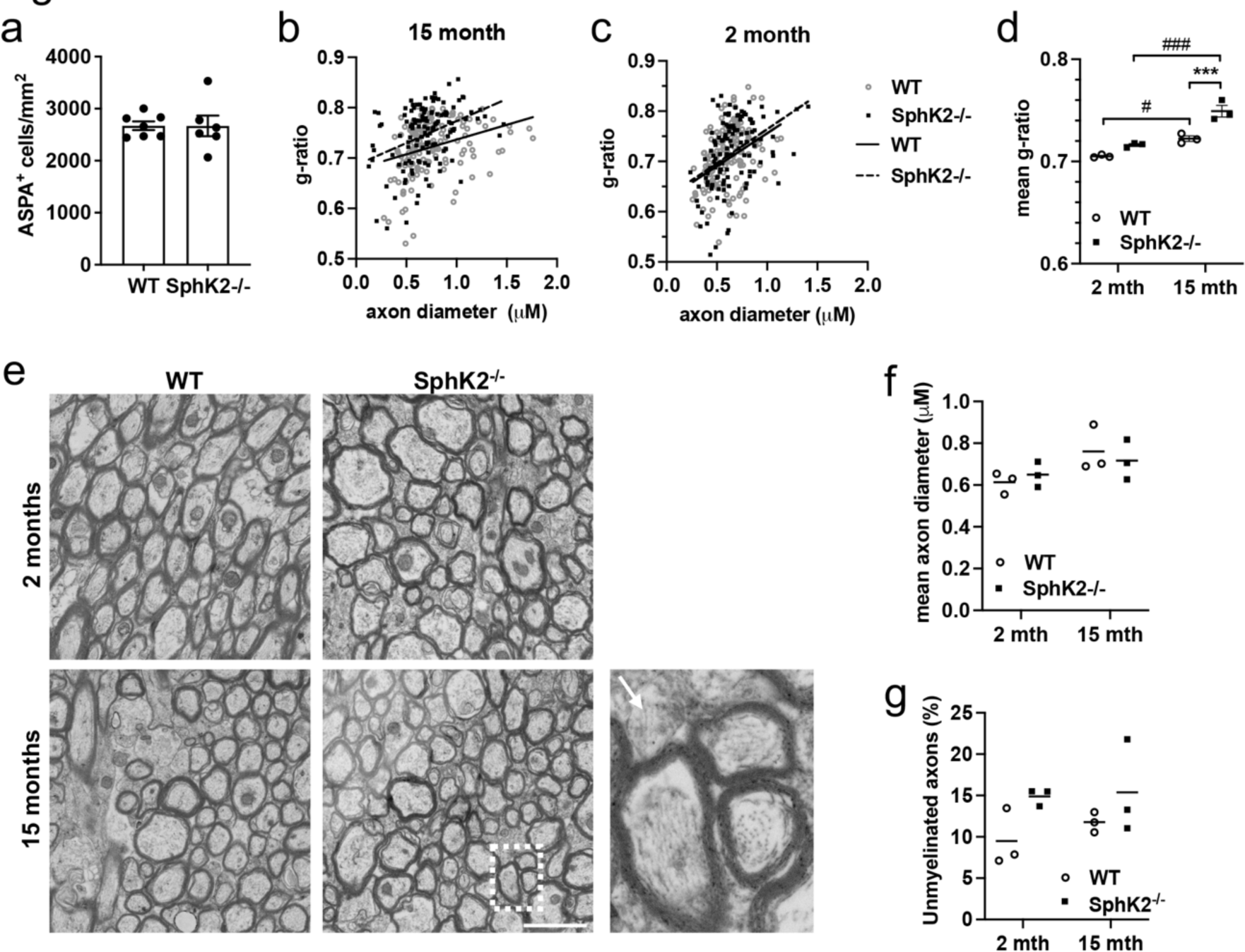
Thinner myelin in aged SphK2*^-/-^* mice. (a) Density of Aspa-positive cells in the CC of 20 month old WT or SphK2^-/-^ mice (6-7 mice/group). (b-c) G-ratio against axon diameter for myelinated axons in the CC of 15 month old (b) or 2 month old (c) mice (measurements from 3 mice/group). Lines of best fit are shown (WT, solid line and open circles; SphK2^-/-^ dashed line and closed squares). (d) Mean g-ratio (3 mice/group); mth = month. (e) Example EM images of myelinated axons in cross-section. Boxed area is enlarged on the right; the arrow marks an unmyelinated axon. Scale bar, 2 μM. (f) Mean axon diameter (3 mice/group). (g) Proportion of unmyelinated axons (3 mice/group). Results for (d), (f), and (g) were analysed by two-way ANOVA with Bonferroni’s post-test (*** *p* < 0.001, WT vs SphK2^-/-^; # *p* < 0.05 and ### *p* < 0.001, 2 month vs 15 month).

## Discussion

Despite considerable interest in the putative oligodendrocyte-protective and pro-myelinating properties of Fingolimod (Alme et al., 2015; Bechet et al., 2020; H. J. Kim et al., 2011; S. Kim et al., 2018; Nystad et al., 2020; J. Zhang et al., 2015), the role for endogenous S1P and sphingosine kinases in myelination has not yet been elucidated. SphK2 is the dominant isoform catalysing S1P synthesis in the CNS (Lei et al., 2017). This study establishes that endogenous SphK2 protects against cuprizone-mediated loss of mature oligodendrocytes and is essential for remyelination following cuprizone intoxication. Myelin protein levels and oligodendrocyte number are normal in the CC of treatment-naïve SphK2^-/-^ mice at 5 months of age, however the loss of mature oligodendrocytes was more severe in SphK2^-/-^ compared to WT mice after six weeks of cuprizone feeding. Remyelination was observed in the CC of WT mice at two weeks after cuprizone withdrawal, whereas remyelination was not observed in the CC of SphK2^-/-^ mice even after a 4-week recovery period. The well-established accumulation of NG2- positive OPCs in cuprizone-treated mice (Gudi et al., 2009; Mason et al., 2000) was unaffected by SphK2 deletion, and mature oligodendrocyte numbers recovered robustly in the CC of both WT and SphK2^-/-^ mice following cuprizone withdrawal, indicating that SphK2 is not necessary for expansion and maturation of OPCs *in vivo* but is necessary for the synthesis of new myelin by mature oligodendrocytes.

Oligodendrocyte apoptosis is reported to commence in the first or second week of cuprizone feeding, leading to notable myelin loss by 4 weeks (Doan et al., 2013; Gudi et al., 2009; Mason et al., 2000). Greater loss of mature oligodendrocytes in SphK2^-/-^ mice with cuprizone feeding is very likely the result of increased apoptosis, although this was not directly measured. An important role for SphK2 in oligodendrocyte survival is supported by our recent work demonstrating that SphK2 deficiency enhances the age-dependent loss of oligodendrocytes in an Alzheimer’s disease mouse model (Lei et al., 2019). S1P is a pro-survival signalling molecule (Grassi et al., 2019; Hannun & Obeid, 2018) and SphK2 generally promotes cell proliferation and survival (Gao & Smith, 2011; Neubauer et al., 2016). This is not true of all cell types and scenarios, for example loss of SphK2 actually protects renal mesangial cells against stress-induced apoptosis (Hofmann et al., 2008). However, SphK2 overexpression in neurons confers resistance to stress through activation of autophagy (Song et al., 2017), whereas global SphK2 deficiency enhances lesion size and functional deficits following cerebral ischemia in mice (Pfeilschifter et al., 2011). It remains to be determined if endogenous SphK2 protects stressed oligodendrocytes *in vitro*.

S1P produced by SphK2 mediates its effects through autocrine or paracrine stimulus of S1P receptors (Choi & Chun, 2013; Grassi et al., 2019), and through intracellular binding targets such as histone deacetylases and peroxisome proliferator-activated receptor γ (PPARγ) (Hait et al., 2009; Parham et al., 2015). Fingolimod-phosphate and S1PR1-selective agonist CYM- 5442 protect oligodendrocytes against apoptosis induced with cuprizone *in vivo* (S. Kim et al., 2018), implying a role for S1PR1 on CNS cells in oligodendrocyte survival. This protection probably results from agonism rather than functional antagonism of S1PR1, since cuprizone-mediated demyelination is significantly worse in mice lacking S1PR1 in oligodendrocytes (H. J. Kim et al., 2011). S1PR5 is very highly expressed by mature oligodendrocytes and mediates pro-survival S1P signalling *in vitro* (Jaillard et al., 2005). No publications have reported on the requirement for S1PR5 in oligodendrocyte protection or remyelination following cuprizone feeding, however S1PR5 deletion impeded remyelination mediated by the S1PR1/5 agonist Siponimod in a demyelination model using *Xenopus* oocytes (Mannioui et al., 2017). Future research should investigate whether pharmacological S1P receptor stimulus rescues oligodendrocyte survival during cuprizone feeding in SphK2^-/-^ mice, and/or restores remyelination following cuprizone withdrawal.

Counting of OPCs and mature oligodendrocytes indicated that OPC expansion and maturation are not affected by SphK2 deficiency. Western blotting showed up-regulation of the oligodendrocyte markers ASPA and CerS2 in both WT and SphK2^-/-^ mice following cuprizone withdrawal. In contrast, the myelin markers MBP, PLP, and CNP increased in WT mice 2 weeks after cuprizone withdrawal but did not recover in SphK2^-/-^ mice even after 4 weeks of cuprizone withdrawal. These results suggest that SphK2 is essential for synthesis of new myelin rather than oligodendrocyte maturation. In accord with this, a recent study using an *ex vivo* nerve injury model demonstrated that S1P and PPARγ are down-regulated during conversion of Schwann cells to a repair phenotype, and up-regulated during re-conversion to a myelinating phenotype (Meyer Zu Reckendorf et al., 2020). Up-regulation of both was necessary for remyelination following nerve injury. S1P stimulates PPARγ-dependent lipogenic gene transcription (Meyer Zu Reckendorf et al., 2020; Parham et al., 2015), which is necessary for myelin synthesis. A limitation of our study is that we did not resolve whether the SphK2 that is required for remyelination is expressed by oligodendrocytes or another cell type. SphK2 deficiency in neurons or other glial cells could produce a lack of S1P receptor stimulus on oligodendrocytes. The related lipid lysophosphatidic acid, produced by dorsal root ganglion neurons, activates Schwann cell LPAR1 receptors to promote their migration (Anliker et al., 2013).

We previously demonstrated that S1P levels are ∼15% of WT levels in the hippocampus and cortex of SphK2^-/-^ mice (Lei et al., 2017; Lei et al., 2019). In this study, we report an equivalent reduction in S1P in the CC of treatment-naïve SphK2^-/-^ mice. Surprisingly, S1P levels were an order of magnitude higher in CC of cuprizone-treated SphK2^-/-^ mice and remained elevated following cuprizone withdrawal. In contrast, S1P levels were modestly reduced in CC of cuprizone-fed WT mice and increased 60% after cuprizone withdrawal. Based on this result, greater loss of mature oligodendrocytes and deficient remyelination in SphK2^-/-^ mice with cuprizone feeding cannot be attributed to bulk S1P deficiency. However, the loss of SphK2- dependent S1P synthesis might still impede oligodendrocyte functions through a deficiency in autocrine, localised S1P receptor stimulus. In this regard, S1PR5 levels were much lower in treatment-naïve SphK2^-/-^ mice compared to their WT littermates, suggesting that basal S1PR5 expression is positively regulated by its own ligand. In contrast, S1PR1 levels were not affected in treatment-naïve SphK2^-/-^ mice, but were suppressed in SphK2^-/-^ mice after cuprizone withdrawal. These disruptions to S1P receptor balance in SphK2^-/-^ mice could affect oligodendrocyte survival and function. Alternatively, direct intracellular targets of SphK2-S1P signalling may be essential for remyelination. For example, S1P produced by nuclear-localised SphK2 directly inhibits type I histone deacetylases in neurons, leading to up-regulation of neurotrophic genes such as brain derived neurotrophic factor, and enhanced hippocampal plasticity (Hait et al., 2014). These pathways cannot be rescued by SphK1, which has no nuclear import signal (Igarashi et al., 2003). It is possible, although unverified, that increased S1P in SphK2^-/-^ mice fed cuprizone is attributed to high SphK1 activity in reactive astrocytes and microglia (Fischer et al., 2011; Nayak et al., 2010). We were not confident quantifying SphK1 protein levels in this study, since the SphK1 antibodies that we tested failed to show loss of the putative SphK1 band in SphK1^-/-^ mouse brain tissue (Supplementary Figure 2).

Pronounced accumulation of the SphK2 substrate sphingosine was observed in cuprizone-fed SphK2^-/-^ mice. Sphingosine remained elevated following cuprizone withdrawal in SphK2^-/-^ mice, as did its precursor ceramide. It is likely that ceramide and sphingosine accumulation in cuprizone-fed SphK2^-/-^ mice results from impaired degradation of myelin sphingolipids. GalCer, sulfatide, and SM are catabolised in lysosomes to sphingosine which, due to its biophysical properties, can escape lysosomes and act as a substrate for new ceramide synthesis in the ER. Complete sphingolipid degradation requires conversion of sphingosine to S1P by SphK1 or SphK2, then irreversible cleavage of S1P by S1P lyase (Hannun & Obeid, 2018).

In direct contrast to S1P, sphingosine is a pro-apoptotic signalling molecule (Kagedal et al., 2001; Woodcock et al., 2010) that activates specific protein kinase C isoforms and inhibits pro-survival signalling by promoting phosphorylation of the abundant scaffolding protein 14-3-3 (Hamaguchi et al., 2003; Woodcock et al., 2010). Ceramide is also a pro-apoptotic lipid that promotes cell death in oligodendrocytes (S. Kim et al., 2012; Plo et al., 1999). Previous reports have demonstrated significantly increased sphingosine and ceramide in cuprizone (S. Kim et al., 2012; Yoo et al., 2020) and experimental autoimmune encephalitis mouse models (Miller et al., 2017). A recent study demonstrated that ceramide resulting from SM hydrolysis impedes remyelination after cuprizone withdrawal (Yoo et al., 2020). The authors proposed that excess ceramides cause oligodendrocyte death and myelin decompaction. Sphingosine and ceramide accumulation might therefore contribute significantly to enhanced loss of mature oligodendrocytes with cuprizone feeding and failure of remyelination in SphK2^-/-^ mice. Alternatively, or in parallel, SphK2 deficiency in cells phagocytosing myelin could block remyelination through impaired catabolism of myelin lipids, as observed in mice lacking the microglial lipoprotein receptor Trem2 (Cantoni et al., 2015; Cignarella et al., 2020).

Our observation of significantly reduced sulfatide, but not HexCer or SM levels, with cuprizone feeding is in agreement with a recent publication (Yoo et al., 2020). Decreased sulfatide is almost certainly a direct consequence of myelin loss. Within the CNS, sulfatide is unique to myelin, whereas SM is produced by all cell types (Schmitt et al., 2015). HexCer comprises, which are positional isomers and indistinguishable in reverse phase LC-MS/MS analyses.

We hypothesized that the remyelination deficit in SphK2^-/-^ mice would affect myelin content and integrity with ageing, as physiological myelin replacement over time is impaired. In agreement with this hypothesis, SphK2^-/-^ mice exhibited substantially increased g-ratios in the CC, indicative of thinner myelin, at 15 months of age. Aspa-positive cell density was equivalent between WT and SphK2^-/-^ mice at 20 months of age, indicating that SphK2 deficiency affects myelin maintenance rather than the survival and/or replacement of oligodendrocytes with ageing. A very modest but statistically significant reduction in myelin thickness was reported in the CC of 3 week old mice lacking S1PR1 in mature oligodendrocytes (Dukala & Soliven, 2016). It would be interesting to determine if this phenotype is more pronounced with ageing, since these mice also displayed increased demyelination with cuprizone (H. J. Kim et al., 2011). Myelin thickness in SphK2^-/-^ mice was not significantly different to WT mice at 2 months of age, implying that developmental myelination is not notably affected by SphK2 deficiency. We speculate that S1P production and lipid flux through SphK1, which is highly expressed during development (Liu et al., 2000), is sufficient for developmental myelination. Alternatively, the requirement for SphK2 in remyelination following cuprizone intoxication may reflect distinct functional roles for SphK1 and SphK2. Unlike developmental myelination, remyelination after a demyelinating insult requires clearance of myelin debris by phagocytic cells (Cantoni et al., 2015). A defect in myelin sphingolipid catabolism resulting from SphK2 deficiency could affect myelin turnover with ageing.

### Conclusions

This study has identified essential roles for SphK2 in protecting against loss of mature oligodendrocytes during cuprizone feeding, and spontaneous remyelination following cuprizone withdrawal. Future work will clarify whether SphK2 promotes remyelination through production of S1P that stimulates S1P receptors on oligodendrocytes, or whether the requirement for SphK2 in sphingolipid catabolism is necessary to clear myelin debris and thereby facilitate remyelination. This research is important for establishing whether S1P receptor signalling is essential for remyelination following injury, and therefore whether agonists of S1PRs, already used in MS therapy, possess remyelinating properties.

## Acknowledgements

This research was funded by project grant APP1100626 and Ideas grant APP2002660 from the National Health and Medical Research Council (NHMRC), Australia (A.S.D.); and MS Research Australia grants 18-0477 (J.D.T. and A.S.D.) and 20-0113 (A.S.D.). We gratefully acknowledge subsidised access to the Sydney Mass Spectrometry and Sydney Microscopy and Microanalysis core facilities, and subsidised mouse housing provided by Laboratory Animal Services, University of Sydney.

---

**Supplementary Figure 1:**
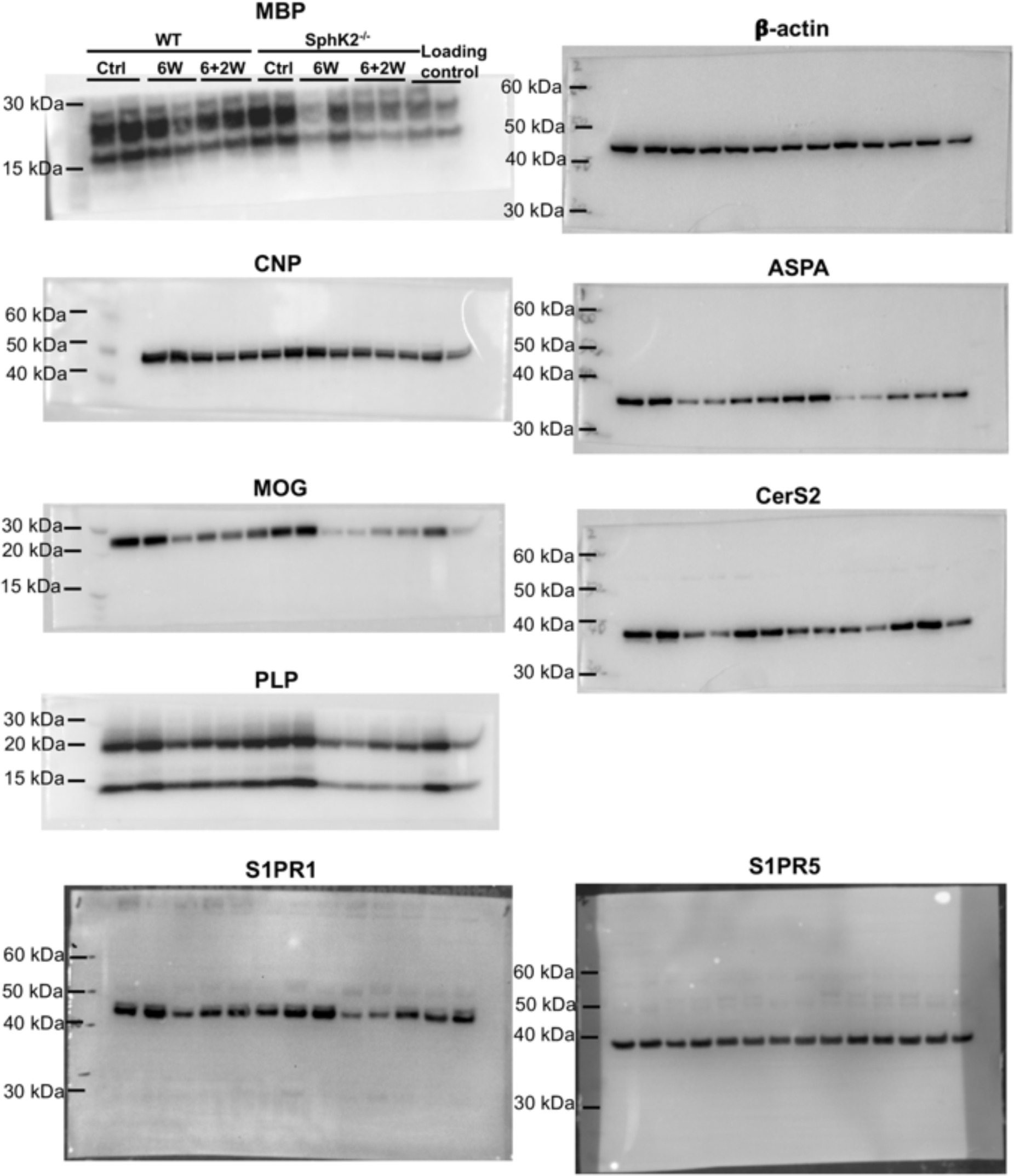
Full images for western blots. Full blot images for antibodies used in this study. Note that for several of these antibodies, the membranes have been cut between 30 and 40 kD to allow blotting for two proteins, so only the higher or lower molecular weight range was probed with a particular antibody.

**Supplementary Figure 2:**
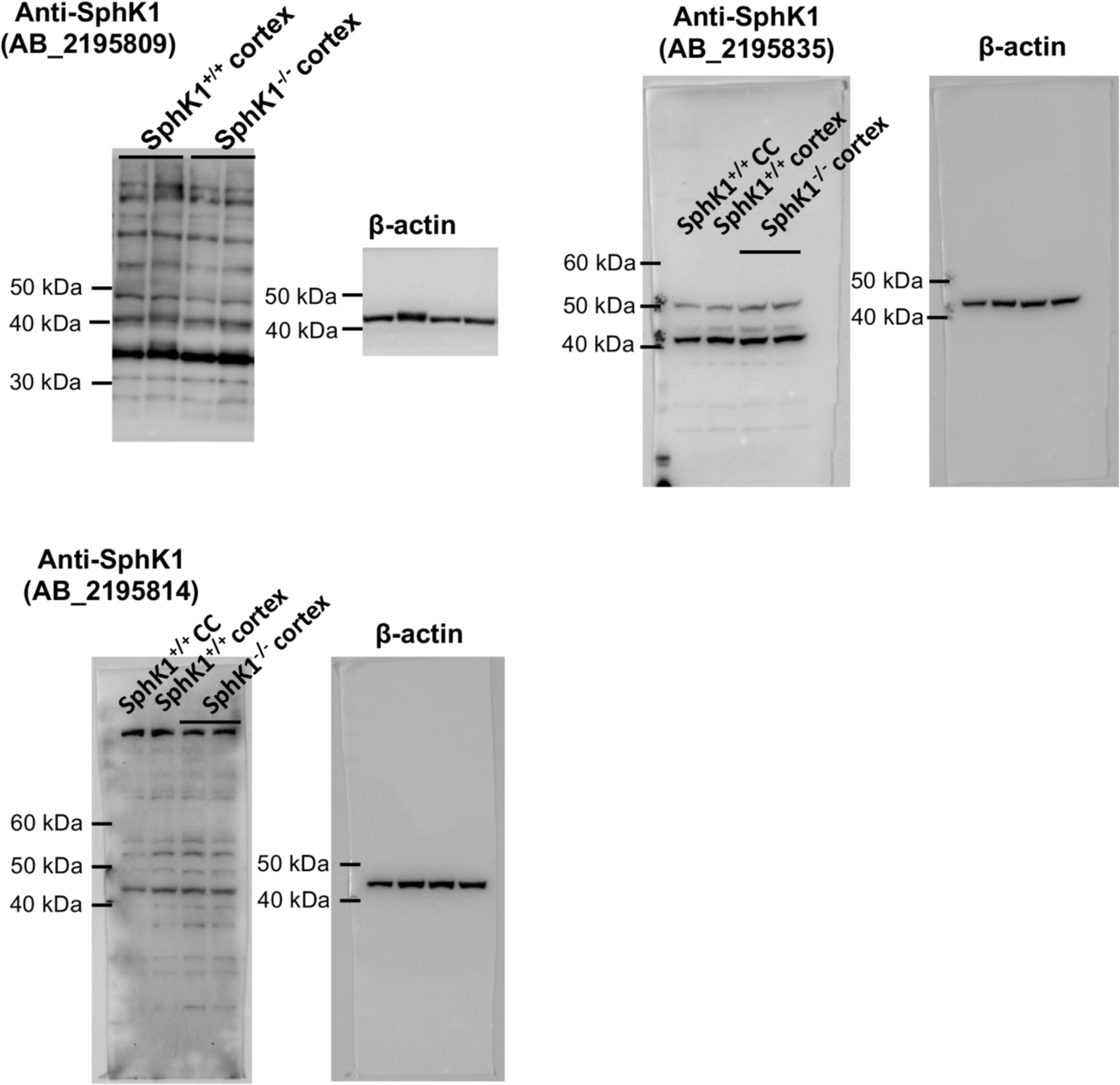
Testing of SphK1 antibodies in mouse brain lysates. Commercial antibodies to SphK1 were tested by western blotting on mouse cortex or corpus callosum (CC) lysates. Specifically, looking for the presence of a band at the correct molecular weight (∼42 kD) in SphK1^+/+^ samples, which is absent in SphK1 knockout (SphK1^-/-^) mouse samples. Membranes were probed with β-actin as a loading control. Antibodies tested were: Proteintech #10670-1-AP, RRID: AB_2195809; Santa Cruz Biotechnology #sc-48825, RRID: AB_2195835 (discontinued); and Abgent #AP7237c, RRID: AB_2195814. Although a major band at the correct estimated molecular weight was observed with antibody sc-48825, this band was also present in lysates of SphK1^-/-^ mice. SphK1^-/-^ mouse samples used for these blots have been described previously (Couttas et al., 2020).

